# Gap junction and amino acid import in somatic cells promote germ cell growth

**DOI:** 10.1101/2023.05.15.540837

**Authors:** Caroline Vachias, Camille Tourlonias, Louis Grelée, Nathalie Gueguen, Yoan Renaud, Parvathy Venugopal, Graziella Richard, Pierre Pouchin, Émilie Brasset, Vincent Mirouse

## Abstract

Gap junctions allow the exchange of small molecules between cells. How this function could be used to promote cell growth is not yet fully understood. During *Drosophila* ovarian follicle development, germ cells, which are surrounded by epithelial somatic cells, undergo massive growth. We found that this growth depends on gap junctions between these cell populations, with a requirement for Innexin4 and Innexin2, in the germ cells and the somatic cells, respectively. Translatomic analyses revealed that somatic cells express enzymes and transporters involved in amino acid metabolism that are absent in germ cells. Among them, we identified an amino acid transporter required for germline growth. Its ectopic expression in the germline can compensate for its absence in somatic cells. Moreover, affecting either gap junctions or amino-acid import in somatic cells induces P-bodies in the germ cells, a feature associated with an arrest of translation. Finally, in somatic cells, innexin2 expression and gap junction assembly are regulated by the insulin receptor/PI3K kinase pathway. Overall, these results support the view that metabolic transfer through gap junction promotes cell growth and illustrate how such a mechanism can be integrated into a developmental programme, coupling growth control by extrinsic systemic signals with the intrinsic coordination between cell populations.

## Introduction

Gap junctions are channels between adjacent cells. Each cells harbour a hemichannel made of six subunits, connexins in vertebrates and innexins in other animal phyla where gap junctions are found. These two protein families are very different in terms of primary sequence, but similar in terms of conformation (1–4). Hemichannels can be homomeric or heteromeric in composition and channels can be formed from two identical or different hemichannels, leading to a large range of channel combinations with different regulations or solute specificities. Gap junctions generally assemble in plaques that contain many channels and each channel allows the direct communication, and thus molecular flows, between the cytoplasm of two cells. However, due to the channel size and conformation, only passive transfer of small molecules and ions, up to approximatively 1KDa, including second messengers (e.g. inositol triphosphate and calcium) is allowed. Gap junction proteins also participate in other cellular mechanisms, such as cell migration (5,6). Moreover, many molecules that can diffuse through gap junctions are linked to energetic metabolism (e.g. glucose and pyruvate) and to anabolic metabolism (e.g. amino acids). However, despite many studies on these metabolite flows, their functional relevance has been elusive.

Such metabolite exchanges could promote cell growth. Available genetic data suggest that the growth of mammalian oocytes could be an illustration of such a mechanism. Indeed, mutation of connexin 37 (*Cx37*) in germ cells or of *Cx43* in somatic granulosa cells, which surround and are in contact with the oocyte, strongly impairs oocyte and follicle growth (7,8). Moreover, amino acid import and pyruvate production are more efficient in follicle cells, and granulosa cells are required for an effective uptake of some amino acids, including alanine and proline, by the oocyte (9–12). These data suggest a potential metabolic flow towards the oocyte via the gap junctions. Nonetheless, it is not known: i) to which extent this putative transfer of metabolites contributes to explain gap junction impact on oocyte growth; and ii) whether such mechanism is coupled with follicle cell growth and more generally, is integrated in the genetic programme controlling follicle development.

A major feature of oocytes is their large size throughout animal evolution, despite important variation among species (13,14). Their size prefigures the early embryo size and their content will determine the early embryo development success rate and quality (15,16). Hence, oocyte growth is an essential step that requires robust underlying mechanisms (17,18). Oocyte development and growth usually occur in ovarian follicles where germ cells are surrounded by follicle cells (i.e. somatic epithelial cells). *Drosophila* oogenesis takes place in a structure called ovariole, in which follicles continuously arise and develop from the anterior to the posterior end. Each ovary is subdivided in about 16 ovarioles (Fig. 1A). Follicles bud from a structure called the germarium that contains germ cells and somatic stem cells. Follicle development is divided in 14 morphological stages during which the oocyte volume increases 1000 times in about 3 days before being fertilized and laid (19). At stage 1, a follicle contains a germline cyst of 16 interconnected cells: 15 nurse cells that will grow by endoreplication and 1 oocyte at the posterior end, blocked in early meiosis. Each germline cyst is encapsulated by the follicular epithelium that is initially composed of about 30 cells. These cells proliferate giving 800 cells at stage 6 and then they undergo endoreplication (20). Both germ cell growth and follicle cell growth are cell autonomously sensitive to the usual growth systemic signals, such as the Insulin Receptor/Phosphatidylinositol 3 Kinase (InR/PI3K) and Target of Rapamycin (TOR) pathways (18,21–23). These pathways affect also non-cell autonomously the growth of the adjacent tissue in both direction: the soma influences germline growth and *vice versa* (18,22–25). The most striking illustration of this reciprocal coordination is observed when *Pten*, a InR/PI3K repressor, is mutated in follicle somatic cells. The faster growth of these somatic cells induces the faster growth of the surrounded wild-type germ cells. Regulation of somatic cell growth by germline growth involves the Hippo pathway and its modulation by tension induced on the epithelium due to the germline cyst volume increase (26). However, it is not known how the follicle somatic cells control germ cell growth.

**Figure 1:**
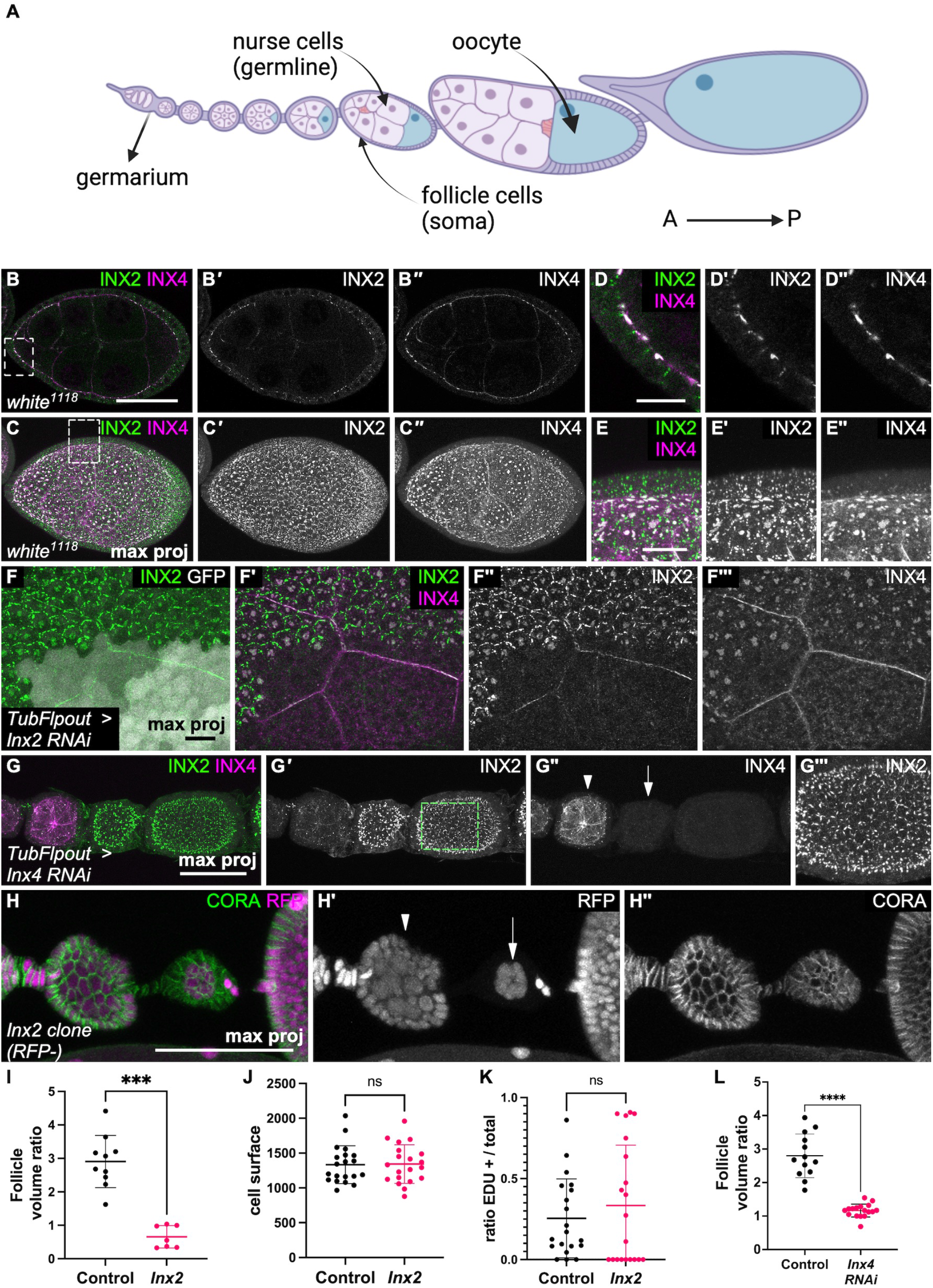
Soma-germline gap junctions are required for germline growth A) Scheme of an ovariole, oriented from the anterior (A) to the posterior (P) end, as all the subsequent images. The ovariole starts with the germarium from which young follicles bud before undergoing massive growth until the formation of an egg at the posterior end. In each follicle, somatic follicle cells (in purple) surround a germline cyst with the nurse cells (pink) and the oocyte (in blue). B) Sagittal view and C) Maximum intensity projection images of a stage 7 follicle after immunostaining for Inx2 (green in B and C, white in B’ and C’) and Inx4 (magenta in B and C, white in B’’ and C’’). D) and E) Higher magnification of the insets in B and C to illustrate the formation of plaques at the germline-soma interface. F) Immunostaining for Inx2 (green in F and F’, white in F’’’) and Inx4 (magenta in F’, white in F’’) in a follicle containing a *Inx2* RNAi-expressing clone in follicle cells marked by GFP expression (white in F). G) Maximum intensity projection of immunostaining for Inx2 (green in G, white in G’’ and G’’’) and Inx4 (magenta in G, white in G’) in follicles containing *Inx4* RNAi-expressing germline clones visualized by Inx4 absence. Note the absence of Inx2 plaques at germline contacts at higher magnification (G’’’) and the growth defect of *Inx4* RNAi follicles (arrow) compared with the wild-type younger follicle (arrowhead) (G’’). H) Somatic *Inx2^A^* mutant clones, marked by the absence of RFP expression, induce a germline growth defect when they cover the whole epithelium (arrow) compared with the wild-type younger follicle (n-1) (arrowhead). I) Quantification of the ratio between the volume of the follicle showing a growth defect fully covered by *Inx2* mutant cells, and the volume of the n-1 follicle of the same ovariole (n= 10 control and 7 *Inx2* mutant follicles, Mann-Whitney test). J-K) quantification of J) cell surface and K) EDU-positive cells in small mutant *Inx2* clones compared with the surrounding wild-type cells (n= 20, 20 clones or groups of wild-type cells for J and K, Welch’s t test). L) Quantification of the volume ratio between a follicle with *Inx4* RNAi in the germline cyst and the wildtype n-1 follicle of the same ovariole (n= 13 for control and n=17 for *Inx4* RNAi, Mann-Whitney test). For all graphs, data are the mean ± SD. ***p <0.01, ****p <0.0001. Scale bars: 50 μm in B,G,H and 10 μm in E,F.

Here, we tested the hypothesis of an evolutionary conservation of gap junction function between germ cells and follicle cells to regulate *Drosophila* female germ cell growth and explored the underlying mechanism. Several reports indicate that gap junctions are present during fly oogenesis. In gem cells, innexin 4 (Inx4) forms plaques at the contact with follicle cells (27–29). Several innexins, including Inx2, are expressed in follicle cells and partially colocalize with or are juxtaposed to Inx4 (28,29). Inx4 is required for germ cell survival before follicle formation, and Inx2 is involved in the germline cyst encapsulation by follicle cells during follicle budding (27,30,31). Inx2 and Inx4 are also required for morphogenetic events during oogenesis. such as the stretching of anterior follicle cells and the migration of border cells during stage 9 (6,29). Moreover, Inx4 and Inx2 form gap junctions in the testis where they are required at different steps of sperm development (27,32,33).

We found that Inx4/Inx2 gap junctions are required for female germline growth once follicles are formed. Moreover, follicle cells specifically express genes involved in amino acid biology, and amino acid import in follicle cells is necessary for germline growth. This germline growth control by somatic follicle cells is linked to the formation of processing bodies (P-bodies) and can be partially bypassed by the direct expression of an amino acid transporter in the germline. Moreover, gap junction assembly is controlled by the InR/PI3K pathway, thus connecting growth coordination between these two cell types with systemic growth control.

## Results

### Inx2 and Inx4 form gap junctions at the soma-germline interface that are required for germline growth

As gap junctions are required for oocyte growth in mammals, we asked whether they could play a similar role in *Drosophila* oogenesis (7,8). Therefore, first, we characterized the gap junction composition and profile in somatic and germ cells, focusing on stages 1 to 8 when follicle development is globally limited to growth without major morphogenetic changes. As previously described (27–29), we detected Inx2 and Inx4 forming plaques at the interface between germ and somatic cells. Inx2 also formed plaques on the lateral domain of follicle cells (Fig. 1B-E). Inx2 expression and plaque size tended to increase as follicles developed (Fig. S1A). The cytoplasmic signal localization suggested that Inx2 and Inx4 were mainly expressed in the soma and germline, respectively. Accordingly, *Inx2* knock-down (*Inx2* RNAi) induction in somatic clones led to the cell-autonomous loss of Inx2 staining, whereas induction of *Inx4* knock-down (*Inx4* RNAi) in the germline led to the loss of Inx4 staining (Fig. 1F,G). Notably, we did not detect Inx4 at the contact of *Inx2* RNAi follicle cells, confirming published results (Fig. 1F’’’) (29). Similarly, when *Inx4* was knocked down in germline cysts, we did not detect Inx2 plaques at the germline contact, although Inx2 protein expression was increased in follicle cells (Fig. 1G’’’). This last observation could suggest a feedback mechanism between gap junction formation and Inx2 expression. Moreover, the loss of *Inx2* in the germline or *Inx4* in the soma did not induce visible defects (Fig. S1B,C). Altogether, these data clearly established the presence of gap junction plaques composed of Inx4 in the germline and of Inx2 in follicle cells, and the Inx4-Inx2 interdependence for plaque assembly. We then tested whether these gap junctions influenced germline growth by generating large *Inx2* mutant clones. In such conditions, we observed follicles that were smaller than the younger one at their anterior, a phenomenon never observed in the wild-type ovarioles. The quantification of the ratio between the volumes of the defective growth follicle and the follicle at its anterior confirmed these observations (Fig. 1H,I). We observed this phenotype with three different alleles, and only when all (or almost) epithelial cells of a follicle harboured the mutated *Inx2* while the anterior one was wild-type or contained a smaller percentage of mutant cells. Similar cases with control clones had no visible effect (Fig. S1D). Because it is known that germline and somatic growth influence each other, we aimed to determine whether *Inx2* effect on germline growth was due to a cell-autonomous effect on somatic growth (22,24,34). We analysed follicle cell growth in smaller clones that do not induce germline defect to avoid feedback between tissues. In such mutant cells we did not observe any difference in cell size and in the proportion of cells in S phase (EDU-positive) between these populations, indicating that Inx2 did not influence somatic growth in a cell autonomous manner (Fig. 1J-K). This suggests that gap junctions are specifically required for germline growth. We could not confirm this hypothesis using germline null *Inx4* mutant clones because in these mutant germline cysts development is blocked very early in the germanium (27). Therefore, we generated *Inx4* RNAi clones in the germline, directly detected by the absence of Inx4 expression (Fig. 1G). *Inx4* RNAi cysts were not larger than the younger wild-type follicle, indicating defective germline growth (Fig. 1G,L). Altogether, these data indicate that Inx2 and Inx4 form homomeric and heterotypic gap junctions between somatic and germ cells that are required for germ cell growth.

### Genes implicated in amino acid metabolism are enriched in somatic follicular cells

Gap junction requirement for germ cell growth supports a model in which metabolites diffuse from follicle cells to germ cells. However, the direct identification of such metabolites is technically challenging. We hypothesized that if follicle cells produce or import metabolites not present in the germline, we might identify enzymes or transporters that are involved in this process and are expressed only in follicle cells. To this aim, we performed translating ribosome affinity purification (TRAP), an approach that allows identifying the tissue-specific translatome. We used *nanos:Gal4VP16* and *trafficjam:Gal4* drivers to express a GFP-tagged ribosomal protein (UASp:RPL10a-GFP) specifically in the germline and in somatic cells, respectively (Fig. 2A). Then, we immunoprecipitated the GFP-tagged polysomes and isolated and sequenced the associated mRNAs under translation. Our biological replicates were highly reproducible (Table S1 and Fig. S2). To extract genes with a specific or strongly enriched somatic expression we used a fold-change of 5 between soma and germline as cut-off that gave a list of 811 genes (Fig. 2B). We then selected genes involved in small molecule metabolic processes, reducing the list to 52 genes. The whole ecdysone synthesis pathway (six genes), which is exclusively active in follicle cells, was enriched in follicle cells, thus validating our TRAP approach (35). Moreover, six genes encoding enzymes involved in amino acid biosynthesis were specifically expressed in follicle cells (Fig. 2B). By analysing this GO class, we found that 29 enzymes were expressed in both cell types, but none was germline-specific (Fig. 2C). Similarly, among the 60 amino acid transporters encoded by the fly genome, 21 were expressed in both tissues, 12 were specifically expressed in the soma, and none in the germline alone (Fig. 2B,D). These data indicated that among the genes specifically expressed in follicle cells there is a strong bias for genes involved in amino acid synthesis or import. Given the importance of these metabolites for growth, they are good candidates to be transferred through gap junctions.

**Figure 2:**
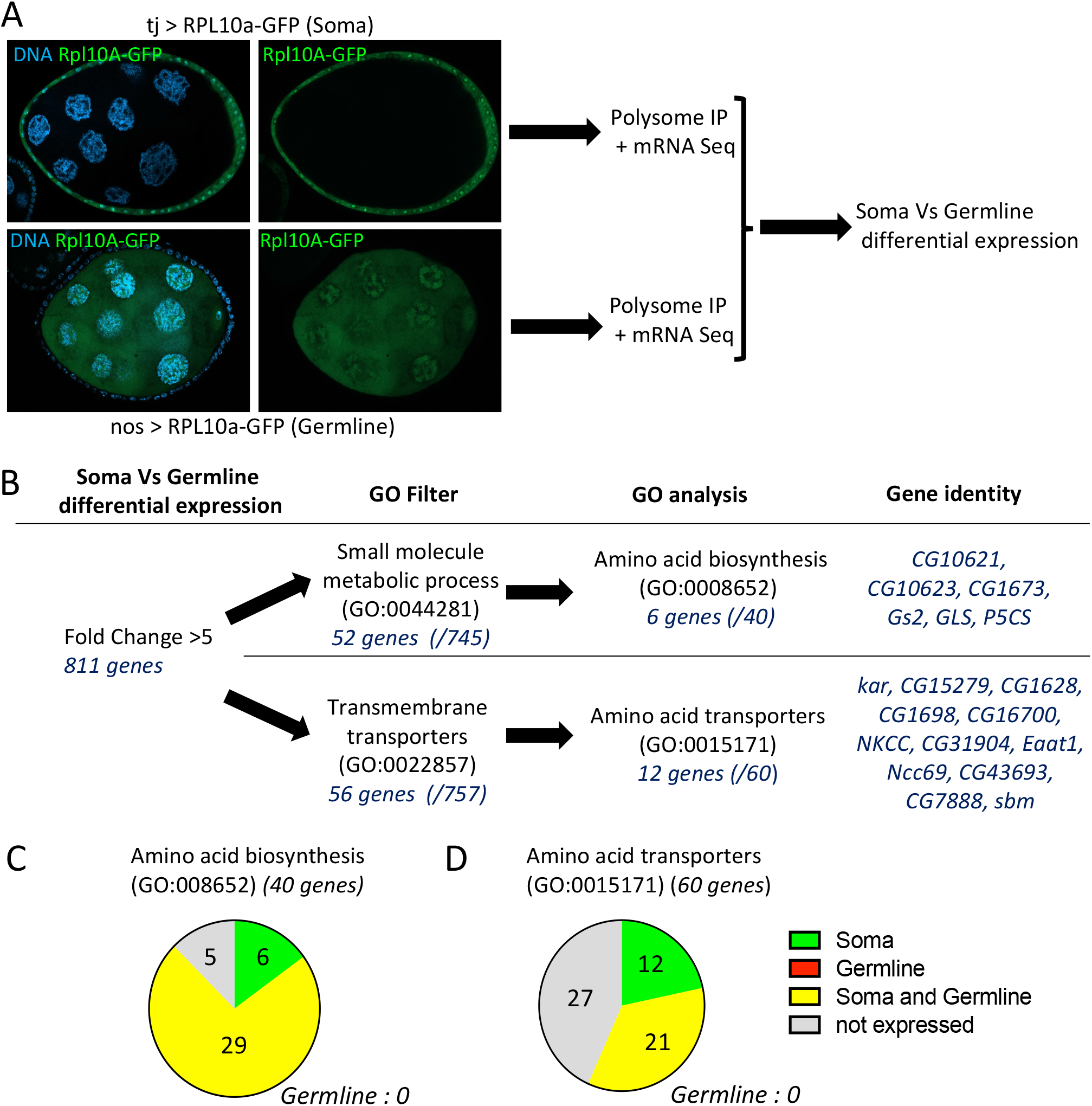
TRAP analysis identifies soma-specific expression of amino acid metabolism and transport genes A) Representative images of follicles expressing RPL10a-GFP in the soma (driven by tj:Gal4) or the germline (driven by nos:Gal4p16). After polysome immunoprecipitation (IP), mRNAs were sequenced. B) The expression of 811 genes was significantly enriched in somatic cells compared with germline cells (fold change >5, p-value <0.05). After filtering for genes involved in small molecule metabolism or transport, genes related to amino acid biology were identified. C) Distribution (soma and/or germline) of genes encoding proteins implicated in amino acid biosynthesis or transport according to their expression profile. No gene of these classes was enriched exclusively in the germline.

### Amino acid import in follicle cells is required for germline growth

Then, we performed a reverse genetic screen by inducing the silencing (RNAi) of the genes involved in amino acid synthesis or transport identified by TRAP in follicle cells and then looking for ovary growth defects. For each amino acid many redundancies may occur between anabolic pathways or transporters that may preclude the observation of clear phenotypes. However, the knock-down of one of the 18 tested genes reproductively induced an ovary growth defect in independent RNAi lines (Fig. 3B-C). This gene, *CG43693*, was uncharacterized in *Drosophila* but belongs to the SLC36 family of amino acid transporters, and was especially close to mammalian SLC36A1 and SLC36A4 that are involved in non-polar amino acids transport as proline, alanine or tryptophan {Boll 2002; Metzner 2006; Pillai 2011

**Figure 3:**
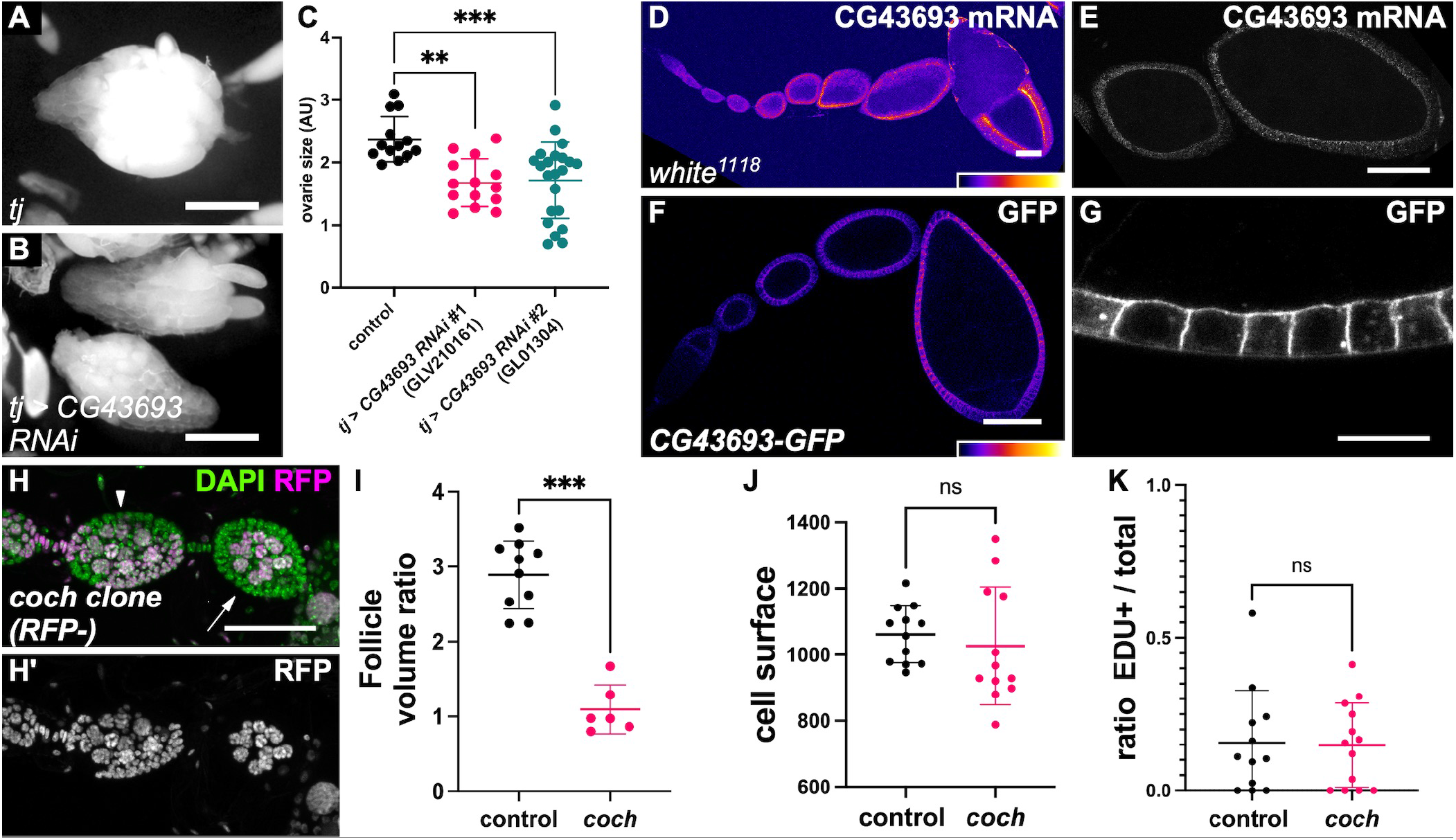
Amino acid import in follicle cells promotes germline growth A-B) Representative images of ovaries in A) control and B) after *CG43693* knock-down by RNAi in somatic cells. C) Ovary size (surface) quantification in control and after *CG43693* somatic knock-down using two RNAi lines (Ordinary one-way ANOVA + Dunnett’s multiple comparisons test). D-E) In situ hybridization analysis of *CG43693* expression in D) a whole ovariole and in E) stage 4 and 7 follicles. F) Expression of endogenous CG43693 protein tagged with GFP from germarium to stage 8. G) Higher magnification of follicle cells showing CG43693-GFP localization in the cell cortex. H) *CG43693/cochonnet (coch)* mutant clones marked by the absence of RFP. When all somatic cells of a follicle are mutated, follicle growth is affected (arrow) as seen when compared with the younger follicle (arrowhead). I) Quantification of the ratio between the volume of follicle with a growth defect and fully covered by mutant cells and the n-1 follicle in the same ovariole (n= 10 control and 6 *coch* mutant follicles, Mann-Whitney test). J-K) quantification of J) cell surface and K) EDU-positive cells in small *coch* mutant clones compared with the surrounding wild-type cells (n=12 control and 12 mutant clones or groups of wild-type cells, Welch’s t test). For all graphs, data are the mean ± SD. **p <0.01, ***p <0.001. Scale bars: 500 μm in A,B, 50 μm D,E,F,H and 10 μm in G.

We confirmed by RNA-FISH that *CG43693* was specifically expressed in follicle cells and checked for its effective knock-down by RNAi in the somatic lineage (Figs 3D-E and S3A). Moreover, we detected *CG43693* mRNA in all follicle cells and at all stages of oogenesis, and its expression progressively increased with the stage (Fig. 3D). A role in importing amino acids from the haemolymph would require a cell membrane localization, while some amino acid transporters are specific to some intracellular vesicular compartments. Therefore, to determine CG43693 protein localization, we generated a GFP knock-in with the MiMIC system to insert in-frame an EGFP-encoding exon (36). This insertion was in the non-conserved N-terminal part common to five of the seven described *CG43693* isoforms. Importantly, it tags isoforms RA and RB that were expressed in follicle cells according to our TRAP mRNA sequence data. The line C*G43693-GFP* was homozygous viable and fertile without visible phenotype. Moreover, ovary size was normal when C*G43693-GFP* was in trans with a deficiency covering the gene, indicating that the insertion did not affect protein function (Fig S3B). We observed a strong GFP signal that tended to increase throughout oogenesis in follicle cells, as previously observed for the mRNA, but no signal in the germline (Fig. 3D,F). The GFP decorates the whole cortex of follicle cells (Fig. 3G). Thus, both CG43693 tissue distribution and subcellular localization were in agreement with the hypothesis that it is implicated in the import of amino acids to promote follicle growth.

We then tested whether CG43693 was required for follicle growth. The Minos element insertion (*CG43693^MI101960^*) used for the GFP knock-in initially contains an exon with STOP codons in different frames. As it is located in the protein N-terminus, it should induce null mutation of the affected isoforms, which include the ones expressed in follicle cells. This allele was sublethal when homozygous or in trans with a deficiency covering the gene. In both cases, the ovaries of these flies were strongly atrophied, fitting with a role for this gene in follicle growth (Fig. S3B). Moreover, Replacing the STOP codon-containing exon with the GFP exon suppressed ovary growth defect. We generated mutant clones in the follicular epithelium. Analysis of small clones that did not cover the whole epithelium did not show any difference in cell size and proliferation, suggesting that follicle cell growth was not affected (Fig. 3J,K). Conversely, when the whole epithelium of a follicle was mutated, such follicles were smaller (or of the same size) than the younger ones, while control clones did not show the same defect (Figs 3H,I and S3C). As this growth defect led to tiny and round follicles, we called this gene *cochonnet (coch)* after the nickname of the small wood ball used with larger metal boules for pétanque, a traditional game in the South of France. Importantly, these experiments indicated that *coch* expression in somatic follicle cells was required for germline growth.

### Genetic evidence of a metabolic transfer between soma and germline cells

Our results support a model in which amino acids imported in follicle cells diffuses through gap junctions to sustain germline growth. In this case, germline expression of *coch* should rescue genetic conditions in which it is absent in somatic follicle cells. Importantly, it is known that the follicular epithelium is permeable at least until stage 8, and thus that metabolites from the haemolymph can directly reach oocyte surface (37). To test this hypothesis, we combined the UAS/Gal4 system with the QUAS/QF system (38). We generated a MatTub:QF driver and a QUASp promoter, inspired by the UASp promoter, for proper expression in the germline (39). A Q*UASp:GFP* transgene driven by MatTub:QF was expressed in the germline at all stages, starting slightly earlier than what described for MatTub:Gal4Vp16, indicating that both driver and QUASp promoter were functional (Fig. S4). Expression of the *QUASp:coch* transgene in the germline with the same driver (*MatTub>coch*) had a slight negative effect on ovary size (Fig. 4A-C,F). However, when it was combined with somatic knock-down of *coch* (*tj>cochRNAi),* it rescued the effect of the latter on ovary size (Fig. 4A,D-F). These genetic data demonstrated that the requirement of *coch* expression in follicular cells for germline growth can be compensated by its germline expression, suggesting a direct metabolic exchange between these cell types.

**Figure 4:**
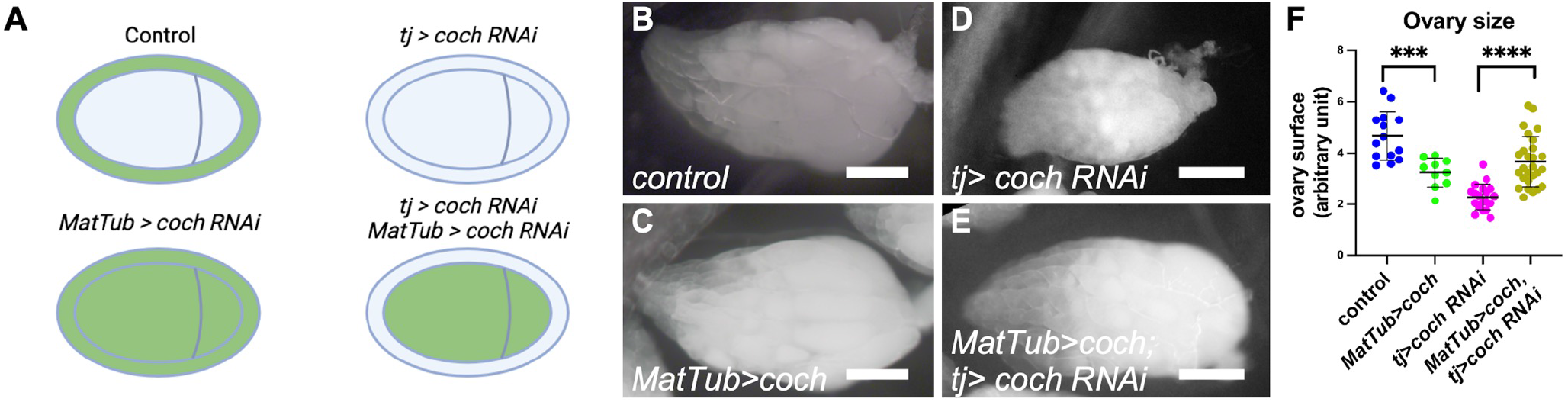
Ectopic amino acid import in the germline compensates its absence in the soma A) scheme representing *coch* expression in green in the different genotypes used on this figure. B-E) Representative images of a full ovary from B) a control female, C) a *MatTub> coch* female D) a *tj > coch RNAi* female, a *MatTub> coch* female and E) a *tj > coch RNAi*, *MatTub> coch* female. F) Quantification of ovary size (surface) in the indicated genotypes (n = 14, 10, 19, and 27 ovaries, Ordinary one-way ANOVA +Tukey’s multiple comparisons test). For all graphs, data are the mean ± SD., ***p <0.001, ****p <0.0001. Scale bars: 500 μm.

### Blocking gap junctions or amino acid import induces processing bodies

Previous studies indicated that somatic growth impairment affects germ cell development (18,22–24). Moreover, the mRNA binding protein Me31B, which is typically concentrated where the anterior-posterior axis determinants localize in the oocyte, becomes enriched in large cytoplasmic structures in the germline when somatic growth is impaired (16,40). These condensates are reminiscent of P-bodies and stress granules observed upon various stresses, including amino acid deprivation (41). In follicles, their formation can be induced by reducing protein availability in fly food and are more easily seen at stage 9 in Me31B-GFP-expressing follicles (40) (Fig. 5A,B). Therefore, we set up a semi-automated method for their quantification by measuring the fluorescence fraction found in condensates in stage 9 nurse cells (Fig. 5C). P-body formation is associated with translation arrest usually induced by inhibitory phosphorylation of the eukaryotic translation initiation factor 2 subunit alpha (eIF2α) on serine 51 (S51) (41). Overexpression in the germline of a transgene mimicking this phosphorylated form (eIF2α-S51D), but not wild-type eIF2α, strongly induced P-body formation, although we did not quantify them because these follicles never reached stage 9 (Fig. 5D,E). Importantly, eIF2α-S51D overexpression was sufficient to completely block germline growth, suggesting a direct link between P-body formation and growth control (Fig. 5E). As P-bodies potentially repress germline growth, we tested whether the absence of gap junctions between germ cells and follicle cells or defective amino acid import in the follicle cells could induce their formation. We knocked down *Inx2* and *coch* in follicle cells using *tj:Gal4* in the presence of Me31B-GFP. Constitutive *Inx2* RNAi in the somatic lineage completely blocked oogenesis and precluded the observation of stages 8-9. Therefore, we added a constitutive *Gal80^ts^* transgene to allow expression of the RNAi construct only during a short period of time at the permissive temperature (14 hours, at 30°C). *Coch* and *Inx2* knock-down strongly induced the formation of P-bodies, suggesting that the observed growth limitation was linked to their formation and the subsequent arrest of translation (Fig 5F-I).

**Figure 5:**
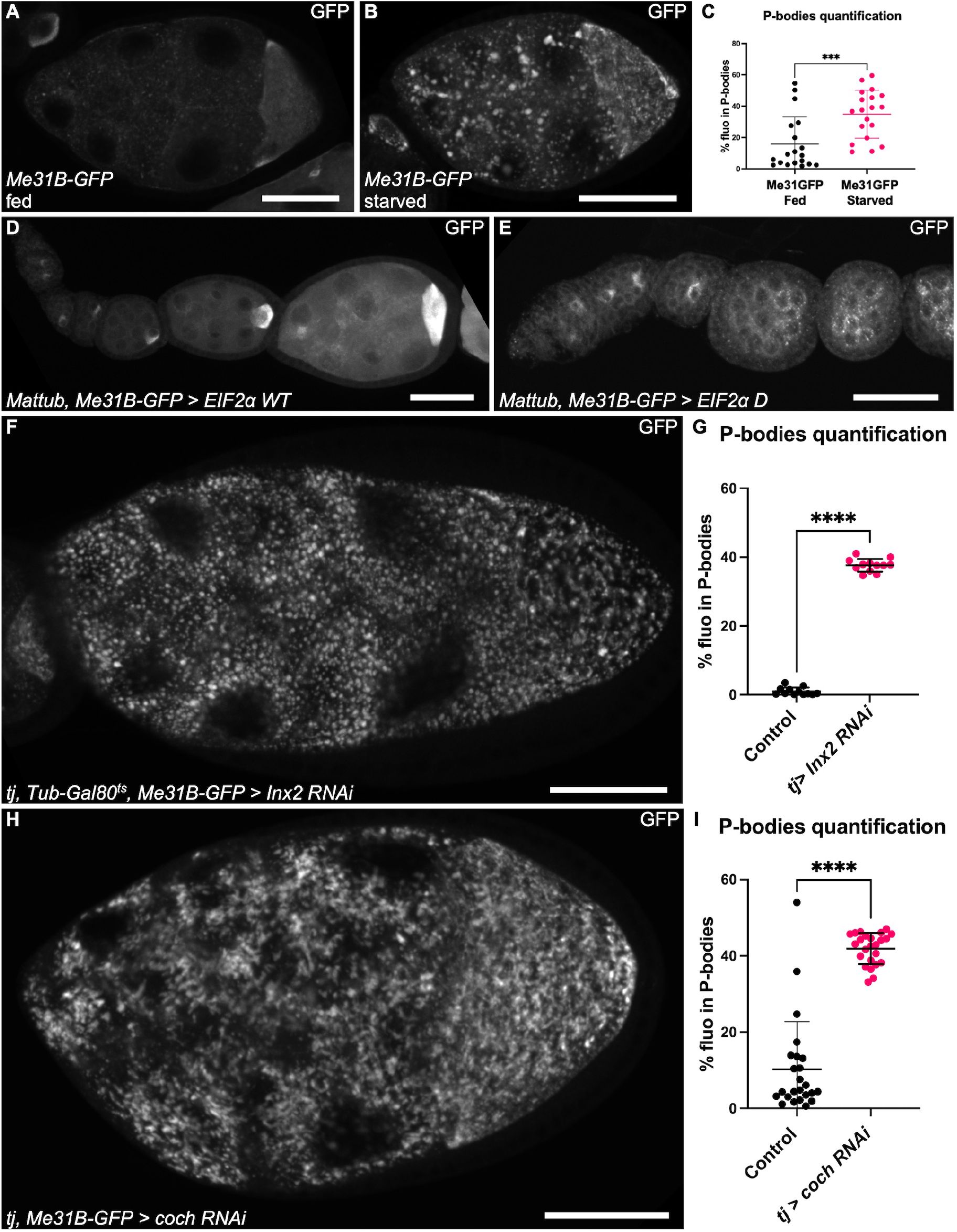
Defective gap junctions or amino acid import induces P-body formation A-B) Endogenous GFP-tagged Me31B protein expression in stage 9 follicles in A) well fed conditions or B) after 15 hours of protein starvation. C) P-body quantification in normal and starved conditions. D-E) Me31B-GFP expression in ovarioles germline that overexpresses D) wild-type eIF2α or E) or eIF2α-S51D (inhibitory phosphorylation). F and H) Representative images of Me31B-GFP expression in stage 9 follicles after RNAi-based knock-down in somatic cells of F) *Inx2* or H) *coch*. G and I) P-body quantification in follicles after RNAi-based knock-down in somatic cells of *Inx2* and I) *coch.* For all graphs, data are the mean ± SD, ***p <0.001, ****p <0.0001, Unpaired t test. Scale bars: 50 μm.

Formation of P-bodies or stress granules following amino acid deprivation can occur independently or dependently of eIF2α S51 phosphorylation. In the first case, which is not the most frequently described, it is associated with the repression of the TOR pathway (42). However, induction of *Tor* RNAi in the germline does not induce P-body formation, excluding this possibility (16). In the second case, eIF2α is phosphorylated by GCN2 that acts as an indirect sensor of amino acid availability. We generated null mutant alleles for *gcn2* that were homozygous viable, as similar alleles described during the course of this project (43–45). Protein deprivation in *gcn2* transheterozygous females still led to P-body formation in the germ cells (Fig. S5A-E). Moreover, an indirect reporter of GCN2 activity and eIF2α−S51 phosphorylation (ATF4-GFP) showed no signal in the germline of wild-type flies, even after protein deprivation (Fig. S5F-G). Lastly, we tested more directly *gcn2* role in the germline when amino acid import in follicle cells is impaired (*tj>cochRNAi*). However, *gcn2* mutant germline clones did not increase the growth rate of wild-type and *coch* RNAi follicles (Fig. S5H,I). These results strongly argued against the implication of GCN2 as a germ cell growth repressor when the availability of the amino acids provided by Coch is reduced.

Altogether these data indicate that although the two well-established pathways involved in amino acid sensing do not seem implicated, P-body formation is linked to germ cell growth control by gap junctions and amino acid import in follicle cells.

### Gap junction assembly links intrinsic growth coordination and systemic control

Published data indicated that InR/PI3K pathway inhibition in follicle cells induces the formation of P-bodies in the germline (16). This result was reproduced by silencing (RNAi) *akt*, an essential actor of this pathway, in follicle cells (Fig. 6A-C). Since somatic activity of the InR/PI3K pathway also strongly influences germline growth, we asked whether there was a link between the InR/PI3K pathway activity in follicle cells and their ability to transfer metabolites to the germline. Coch-GFP expression level and localization were similar in wild-type and in follicle cells with a PI3K gain of function or mutated for *akt,* suggesting no impact on this specific actor of amino acid import in follicle cells (Fig. S6A,B). Conversely, in *akt* mutant cells, Inx2 was almost undetectable (Fig. 6D), whereas Inx2 expression was strongly increased in *Pten* mutant cells and plaques size was increased (Fig. 6E). These data indicated that Inx2 expression is sensitive to InR/PI3K pathway gain and loss of function. We also observed that *Inx2* mRNA level was strongly increased in *Pten* mutant follicle cells, suggesting that the observed effects on protein level and plaque assembly were due *Inx2* upregulation (Fig. 6F). These observations supported a model in which the non-cell autonomous effect of the InR/PI3K pathway from somatic cells to germ cells is mediated by gap junctions. To test this hypothesis, we performed an epistasis experiment. As previously described (18), large *Pten* mutant clones in the follicular epithelium accelerated germline growth in a non-cell autonomous manner (Fig. 6G). This effect was abrogated upon *Inx2* RNAi induction in *Pten* mutant cells (Fig. 6H). Thus, gap junctions are required for germline growth control via InR/PI3K activity in the follicular epithelium. Altogether, our data link the systemic control of somatic cells to the growth coordination between somatic and germline cells via the modulation of gap junction assembly.

**Figure 6:**
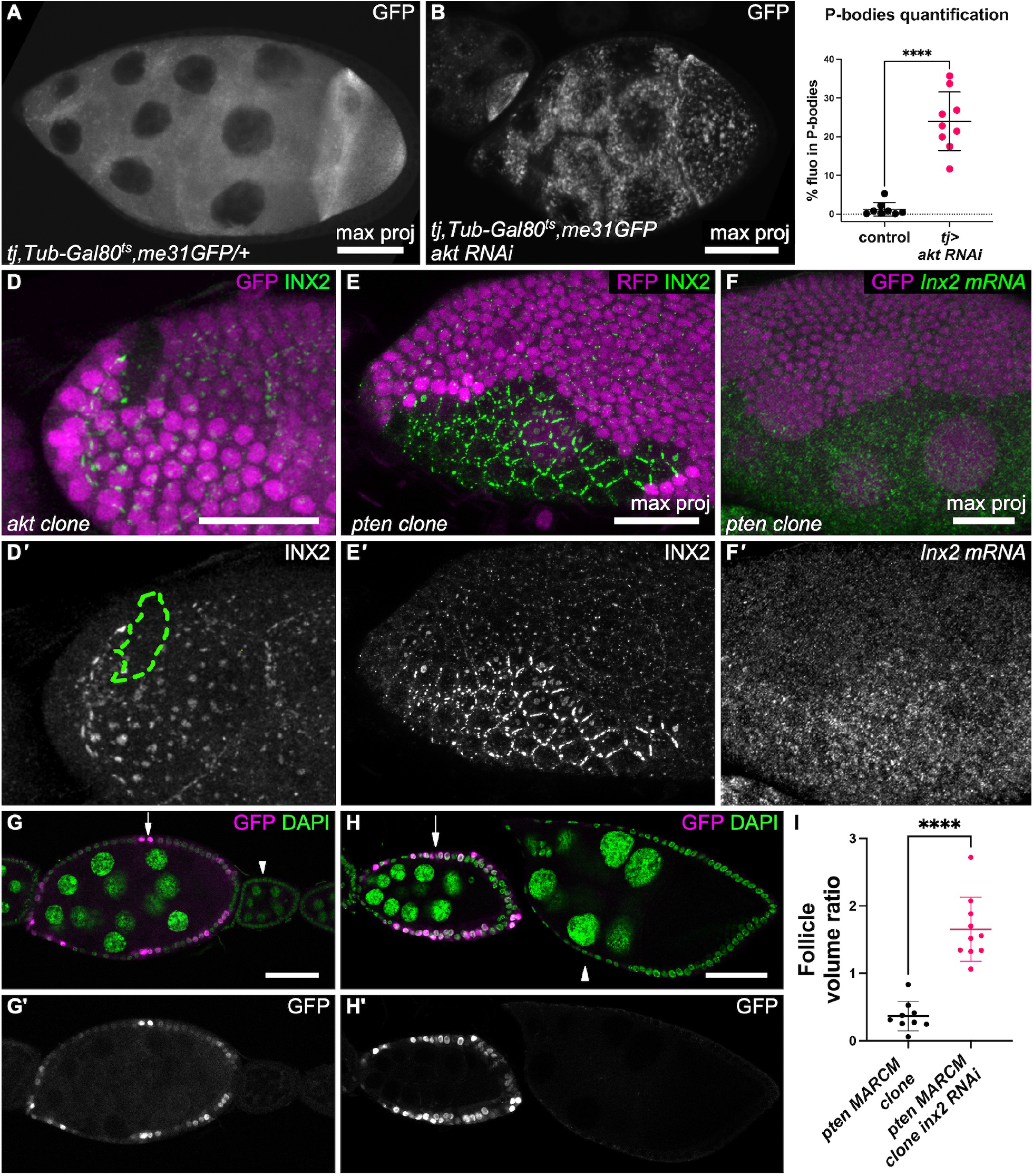
Gap junctions are controlled by the InR/PI3K pathway A,B) Representative images of Me31B-BFP expression in stage 9 follicles from A) a control female and B) a *tj:Gal4, Tub:Gal80^ts^ > akt RNAi* female. C) P-bodies quantification (fluorescence intensity) in the indicated genotypes (n = 8 and 9 follicles, unpaired t test). Data are the mean ± SD. ****p <0.0001. D) Inx2 expression in an *akt^q^* mutant clone (marked by the absence of GFP expression). E-F) Inx2 protein (E) and *Inx2* mRNA expression by FISH (F) in a *Pten^dj189^* mutant clone marked by the absence of RFP expression. G) Large MARCM *Pten^dj189^* mutant clone marked by the presence of GFP expression showing faster follicle growth. H) Large MARCM *Pten^dj189^* mutant clone in which faster follicle growth was abolished after expression of an RNAi against *Inx2*. I) Quantification of volume ratio between the older fully wild-type follicle and follicles containing a majority of *Pten* mutant cells and expressing or not RNAi against *Inx2* and (n = 10 and 9, Mann-Whitney test). Scale bars: 50 μm in A, B, G, H and 20 μm in D,E,F. Data are the mean ± SD. ****p <0.0001.

## Discussion

In this article we show that gap junctions participate in the control of cell growth, leading to the intuitive proposal that they allow a metabolic flow between cells. In accordance with this hypothesis, we identified an amino acid transporter, *coch*, specifically expressed in follicle cells and required for germline cell growth. Though a flux of amino acids from somatic cells to germ cells cannot be directly visualized, *coch* expression in the germline can partially bypass its silencing in the soma, providing evidence for a functional metabolic exchange between these cell types. Moreover, both defective gap junction or amino-acid import in somatic cells induces P-bodies in the germ cells, a feature associated with an arrest of translation. Thus, collectively, our data lead to a model in which gap junctions allow the transfer from somatic cells to germ cells of metabolites, such as amino acids, that germ cells cannot directly produce or import, thereby promoting germ cell growth through translation control.

Inx2 and Inx4 have various functions during *Drosophila* oogenesis (6,27,29,31). However, our data showing their involvement in germ cell growth can be more easily compared with gap junction role in mammal follicles where Cx37 is expressed in the oocyte and Cx43 in follicle cells (7,8). This indicates that gap junction requirement for oocyte growth is a conserved feature throughout evolution. Gap junctions are present in most animal tissues and can directly connect different cell types, for instance, neurons and glial cells. Therefore, their involvement in growth control might be more general. We found that Inx2 expression and the subsequent formation of plaques were strongly regulated by the InR/PI3K pathway in a cell autonomous manner. This indicates that gap junctions are a permissive prerequisite for germline growth, and it also provides an effective mechanism to link metabolic flow with growth systemic signals and coordinate the growth of the two connected cell populations.

Our results also established that Coch, an amino acid transporter of the SLC36A family, must be expressed in somatic cells for germline growth. Amino acid availability and cell growth control are usually linked by the TOR pathway (46). However, this pathway is unlikely implicated in the mechanism described here. Indeed, first, when germline growth is blocked due to alteration of somatic cell growth upon loss of *akt*, which also regulates gap junction assembly, the TOR pathway is still active in the germline (18). Moreover, overactivation of the TOR pathway in the germline in such conditions does not suppress growth inhibition. Finally, TOR inhibition in the *Drosophila* female germline does not induce P-body formation (16). Therefore, we tested the involvement of the other well-described amino acid sensing pathway that relies on GCN2. Our results also excluded this mechanism to explain P-body formation and growth inhibition in the absence of proline import in somatic cells or of gap junctions. Therefore, more studies are needed to determine the precise mechanism underlying germline growth control. Nevertheless, several arguments suggest that amino acids imported by Coch might just be the tip of the iceberg of the metabolic cooperativity between follicle cells and germ cells. *Inx2* mutant clones have a stronger impact on germline growth than *coch* mutant clones, suggesting that other metabolites from follicle cells also may promote germline growth. Accordingly, our translatome analysis showed that different genes involved in amino acid synthesis or import were specifically expressed in follicle cells, but none in germ cells. The absence of phenotype when these genes were knocked down, except *coch*, could be explained by multiple potential levels of redundancy among synthesis pathways, transporters, and synthesis and import. Nonetheless, detailed analysis of the genes overexpressed in follicle cells compared with germ cells suggest that several amino acids (e.g. proline, tryptophane, valine, tyrosine) could flow from the soma to the germline. Moreover, in the mouse, the alanine transporter SLC38A3 is strongly enriched in granulosa cells that are required for efficient alanine import in the oocyte (11). Thus, altogether these data suggest that gap junction-dependent amino acid flow is not restricted to the amino acids imported via Coch. Besides, glucose intake and pyruvate production are more efficient in granulosa cells, suggesting that follicle cells could provide energetic molecules to the oocyte (12). In line with this observation in mammals, TRAP data indicated that transporter for trehalose (tret1-1), the circulating sugar in insects, was expressed in somatic cells and not in the germline, whereas germline development is highly dependent on sugar and the pentose phosphate pathway (47). Thus, a contribution of energetic metabolism in the soma-germline cooperativity via gap junctions will be an interesting avenue for future investigations.

One might ask what is the evolutive advantage of this metabolic cooperativity compared with direct import or synthesis in the germline. Three main hypotheses could be proposed. First, metabolites might not be able to directly reach oocyte membrane due to the presence of the follicular epithelium. However, in follicles from stage 1 to 8, septate junctions are immature, and the epithelium is not impermeable (37). Accordingly, our results showing a rescue of *coch* absence in the soma by its ectopic expression in the germline imply that its solutes can reach the oocyte. Second, female germ cells undergo massive growth and there is a non-linear relation between volume and cell surface increases. Consequently, surface exchange with the extracellular medium might not allow sufficient metabolic import and may require the support of follicle cells. This hypothesis was proposed to explain gap junction requirement for mammal oocyte growth (48). However, in this case, both tissues should be able to import the required metabolites and our data does not support such a model because follicle cells expressed a whole set of anabolic enzymes and transporters that were not expressed in the germline. Alternatively, such a mechanism may provide a protective effect for the germline. It is well established that the cell metabolic activity can induce stress, with for instance, the production of reactive oxygen species (ROS). However, the germline must be protected, especially its DNA content that is transmitted to the embryo. A recent study demonstrated that mammalian oocytes block ROS production by suppressing mitochondrial complex I (49). Thus, a gap junction-mediated metabolic exchange might allow externalizing the stress-generating metabolic activity to follicle cells that will anyway die few days later. Notably, *coch* expression in germ cells had a slight but significant negative impact on ovary size, suggesting that its presence in germ cells is detrimental for follicle development. Although, the exact reason for this deleterious effect is unknown, this observation fits with a model in which externalization of metabolic import and activity could facilitate proper oocyte development. However, Coch is a transporter, and not an enzyme, and thus, the protective effect might not be directly due to amino acid exclusion, but to one of the many possible downstream metabolic activities.

Altogether, our data indicate that gap junctions and a metabolic flow are essential for cell growth and that gap junction assembly can be modulated to adjust cell growth rate. Moreover, it shows that amino acids, and potentially other metabolites, are important actors in this mechanism ending with the formation of P-bodies, opening a large field for investigation to obtain a comprehensive view of this metabolic cooperativity.

## Materials and Methods

### Fly genetics and handling

All fly stocks used are detailed in Table S2. The final genotypes, temperature and heat-shock conditions are in Table S3. *gcn2* null alleles were generated by inducing indel mutations using an available gRNA line. Alleles are described in Table S2. Unless specified, flies were kept on a corn meal-based medium with 80 g/L fresh yeast. Protein starvation was performed on grape juice agar plate. Coch-EGFP was obtained by inserting the EGFP-FlAsH-StrepII-TEV-3xFlag cassette in the Minos element insertion MI01960, as previously described (36).

### TRAP experiments

Ovaries of 100 females of each genotype (tj>RPL10-GFP or Nos>RPL10-GFP) were dissected on ice. Then, TRAP was performed as described in Bertin et al, 2015. Briefly, after homogenization, ovary extracts were preabsorbed with magnetic beads. Immunoprecipitation was performed with anti-GFP antibodies already coupled with magnetic beads before mRNA extraction. Tissue specificity and enrichment of the extracted mRNAs were checked by RT-qPCR with *GFP* (enrichment), *traffic-jam* (soma specific), *Ago3* and *Aubergine* (germline specific) primers. mRNA libraries were made with the Nugen Ovation 1-16 droso Universal RNA-seq kit according to the manufacturer’s instructions. Sequencing was performed by Fasteris. Data were deposited on GEO (GSE230452) and described in Table S1.

### Molecular cloning and transgenesis

The *QF* sequence was amplified from the pAttB-QF-sv40 vector and cloned in the vector that contains the alpha4-tubulin promoter (MatTub). The pQUASp vector was constructed from pUAST in which the promoter was replaced by QUAS sites and the minimal P-element promoter was amplified from the pUASp vector. Then, this vector was used to clone *EGFP* or *coch* coding sequence and P-element insertions were generated. *eIF2α* and *eIF2α-S51D* coding sequences were cloned in the pUASz vector, and transgenes were inserted at the AttP40 landing site. All vectors and new *Drosophila* lines can be provided upon request.

### Immunostaining, FISH, EDU incorporation, imaging and quantitative analyses

Resources and reagents are listed in Table S1. Immunostaining was performed as described in Vachias et al, 2014. Stellaris SM-FISH oligo-probes against *Inx2* and *CG43693/coch* mRNAs were produced by Biosearch Technologies. FISH was performed according to the manufacturer’s instruction. Images were acquired on a Zeiss LSM800 confocal microscope or a Zeiss Cell observer spinning disc microscope. P-bodies were quantified using a homemade macro initially designed to quantify basement membrane fibrils (50). Quantifications were done on 5 μm z-stack projections of spinning-disc images acquired with a x20 lens. After manual selection of a ROI corresponding to nurse cells, P-bodies were detected by keeping objects with a minimal size of 5 adjacent pixels and a minimal fluorescence intensity of x1.75 of the mean intensity. Then, the total fluorescence contained in these P-bodies was quantified and reported as a fraction of the total fluorescence of the ROI. For EDU incorporation, ovaries were incubated with 10μM EDU in complemented Schneider medium for 15 minutes. After fixation, staining was performed according to the manufacturer’s instructions (Kit EDU C10638, Thermo Fisher). Mutant-control follicle volume ratios were calculated after estimating the follicle volume by multiplying their length by their width to the square. Mutant and control follicles in position 2 to 4 of the ovariole starting from the anterior were analyzed. Cell size was automatically determined after cell segmentation using Tissue Analyzer (51). Statistical analyses were performed with Prism. For all experiments, the minimum sample size is indicated in the figure legends. For each experiment, multiple females were dissected. Randomization or blinding was not performed. The sample normality was calculated using the D’Agostino and Pearson normality test. The unpaired *t*-test was used to compare samples with normal distribution (or one-way ANOVA for multiple comparisons), and the unpaired Mann–Whitney test for samples without normal distribution. Figures were prepared using ScientiFig (52).

## Acknowledgements

We are grateful to G. Junion, P Phelan, G Tanentzapf, H.D. Ryoo for sharing antibodies or fly stocks. We also thank the CLIC facility (Clermont Imagerie Confocale). This research was financed by the French government IDEX-ISITE initiative 16-IDEX-0001 (CAP 20-25).

## Supplemental data

**Figure S1.**
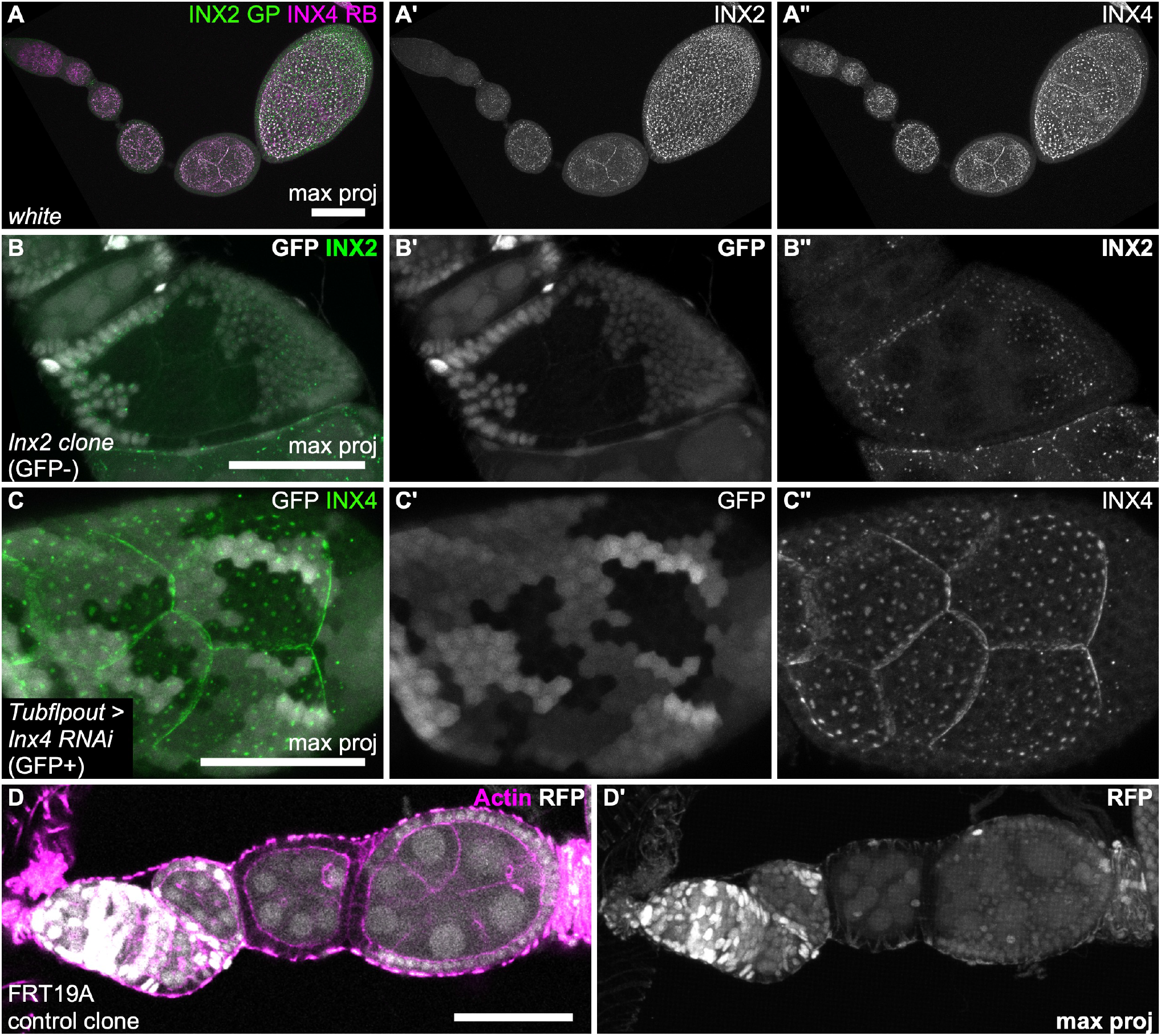
A) Maximum intensity projection images of an ovariole after immunostaining for Inx2 (green in A, white in A’) and Inx4 (magenta in A, white in A’’). Note the progressive increase of Inx2 expression. B) follicle containing a germline mutant clone for *Inx2* (absence of GFP white in B and B’) and showing no growth defect and no impact on gap junction plaques while Inx2 staining (green in B, white in B’’)is lost in the somatic clone observed on the same follicle C) Somatic *Inx4* RNAi clone showing no impact on plaque formation (Inx4 staining green in C,white in C’’). D) Control FRT19A RFP minus clone covering the whole epithelium of a follicle and showing no growth defect when compared to surrounding follicles. Scale bars 50 μm

**Figure S2.**
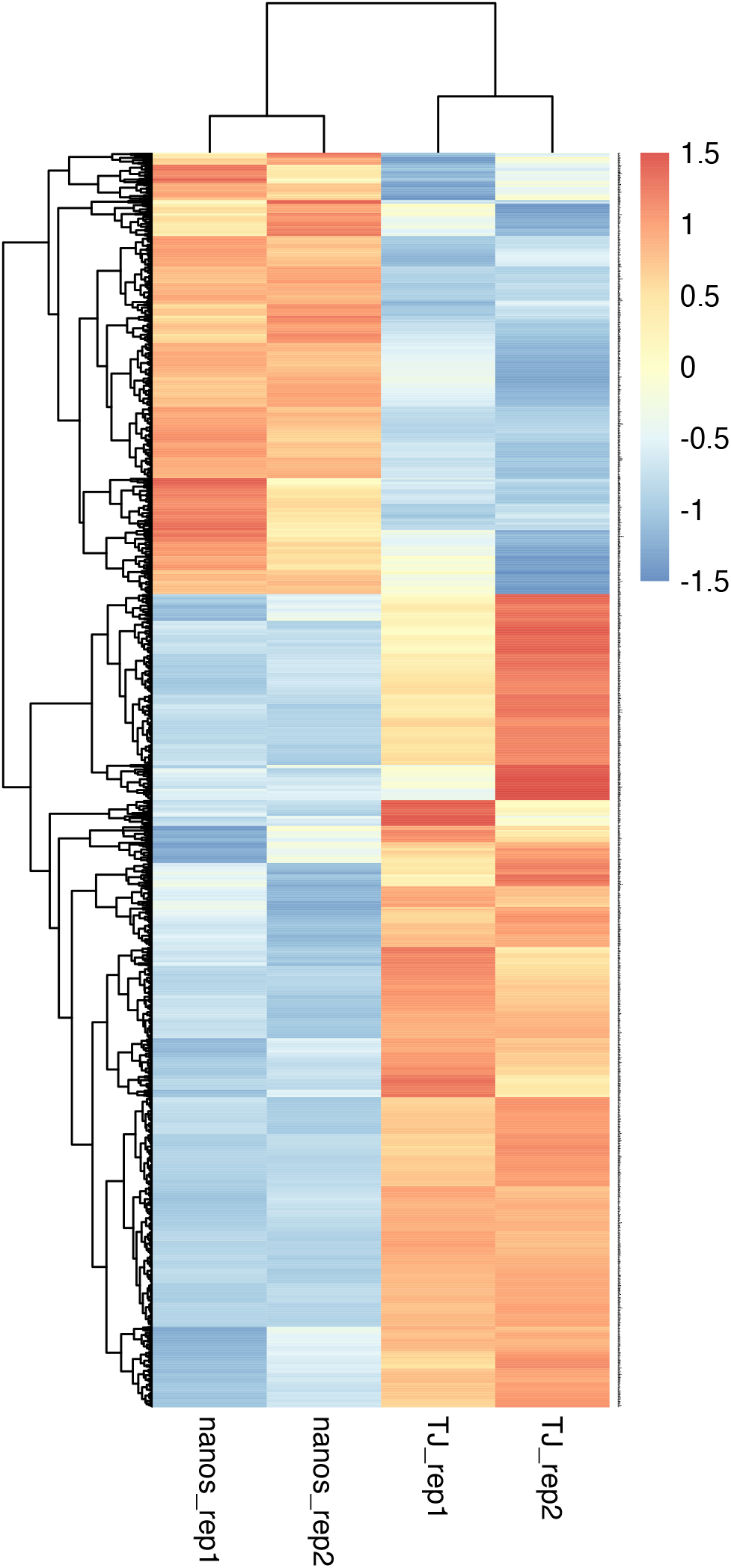
Z-score hierarchical clustering heat map visualization. Only significantly differentially expressed genes are reported. Highly expression correlation can be noticed across condition replicates.

**Figure S3.**
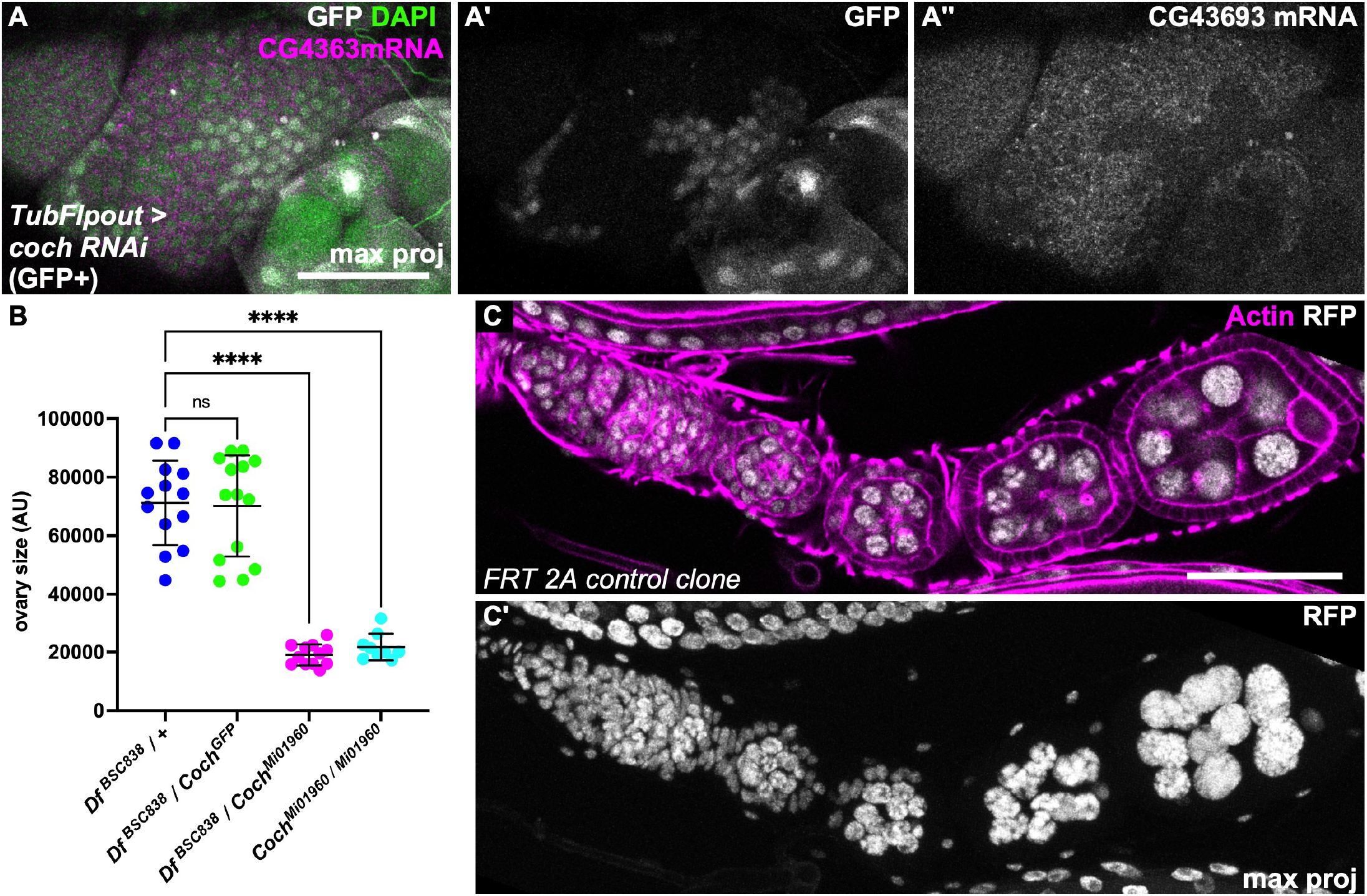
A) RNAi clone against *CG43693/coch* stained by FISH against *CG43693/coch* showing its disappearance from these cells, confirming probe specificity and RNAi efficiency. B) Quantification of ovary size (in arbitrary units) for the indicated genotypes. Data are the mean ± SD. ****p <0.0001. C) Control FRT 2A RFP minus clone covering the whole epithelium of a follicle and showing no growth defect when compared to surrounding follicles. Scale bars 50 μm

**Figure S4.**
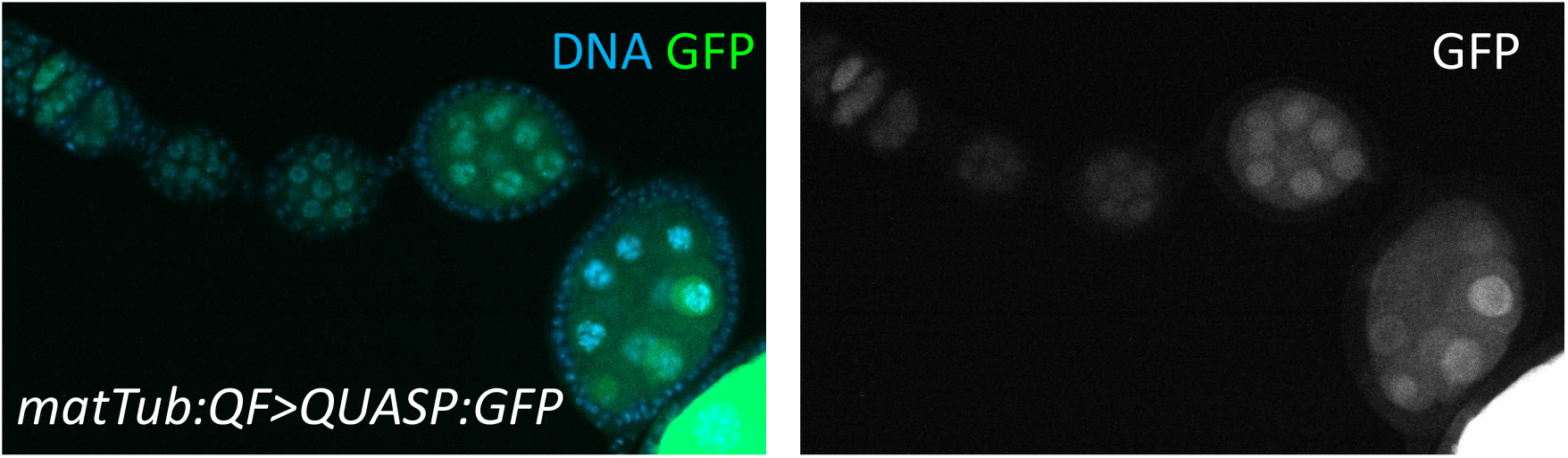
Ovariole expressing a QUASP:GFP transgene under the control of the MatTub:QF driver.

**Figure S5.**
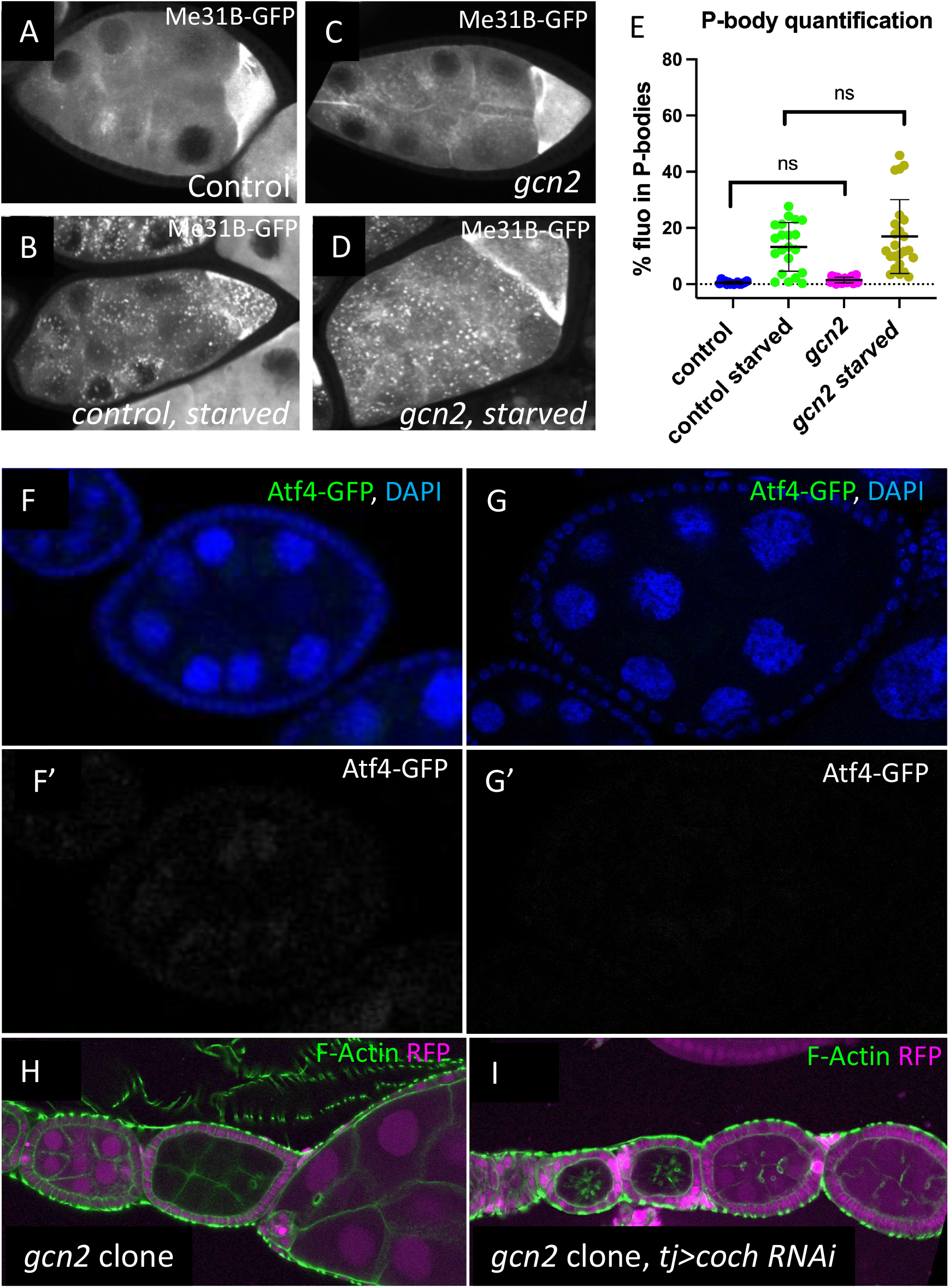
A,B) Representative images of Me31B-BFP expression in stage 9 follicles from a: A) control female, B) starved control female, C) *gcn2* transheterozygous mutant female, and D protein starved *gcn2* transheterozygous mutant female. E) P-bodies quantification (fluorescence intensity) in follicles of the indicated genotypes and conditions. Data are the mean ± SD. F,G) Absence of Atf4-GFP protein expression used as a read-out of GCN2 activity in F) normal and G) protein-starved conditions (green in F and G., white in F’ and G’). H-I) Ovarioles with germline *gcn2* mutant clones marked by the absence of RFP expression (magenta) and stained for F-actin (green) in H) wild-type background and I) in flies harbouring *coch* RNAi in follicle cells.

**Figure S6.**
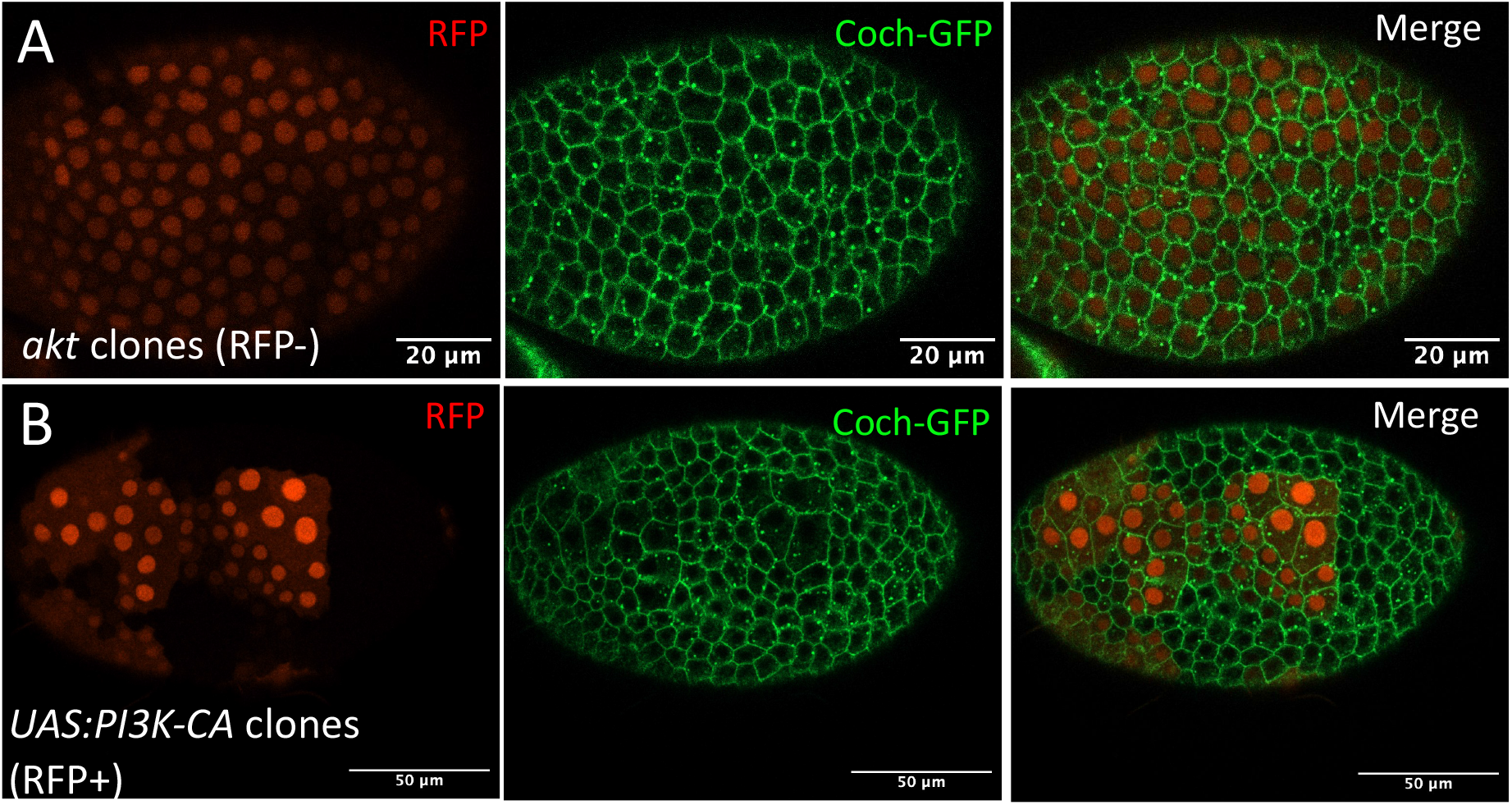
A) Coch-GFP expressing follicle with *akt* mutant clones (RFP-negative cells). B) Coch-GFP expressing follicle with flip-out clones that express a constitutively active form of PI3K (RFP-positive cells).

**Table S1: Excel file of soma vs germline TRAP analysis**

**Table S2:**
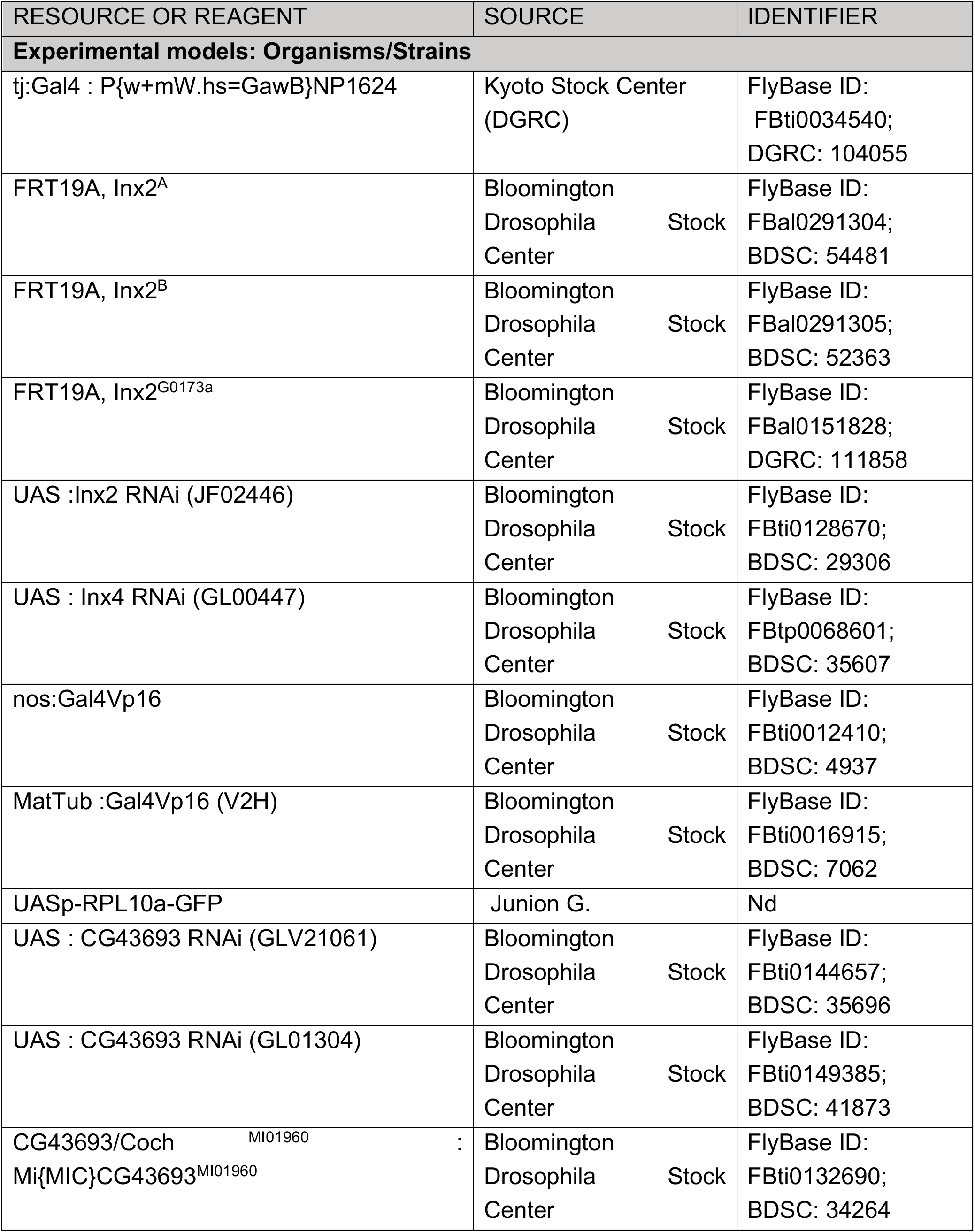
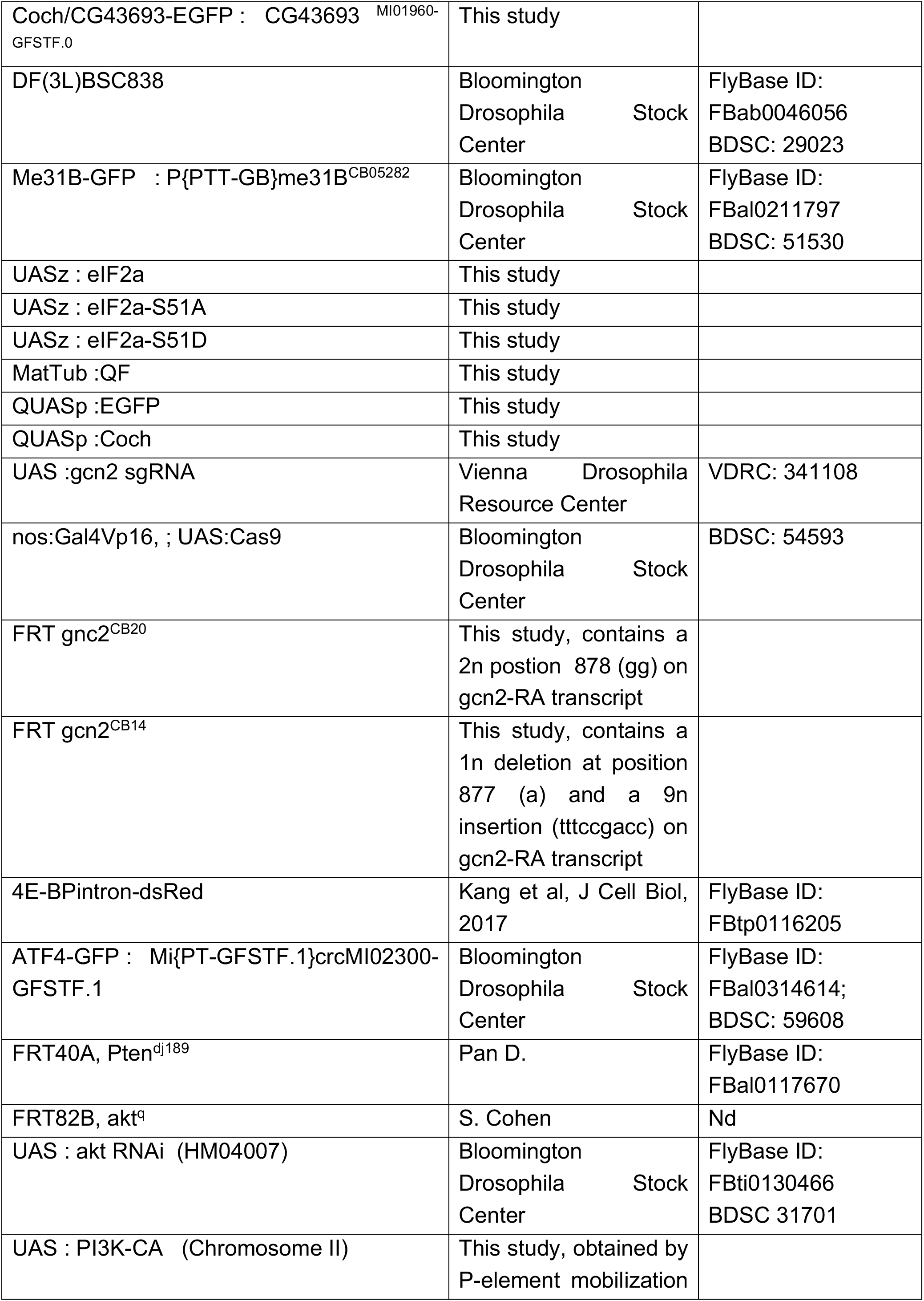
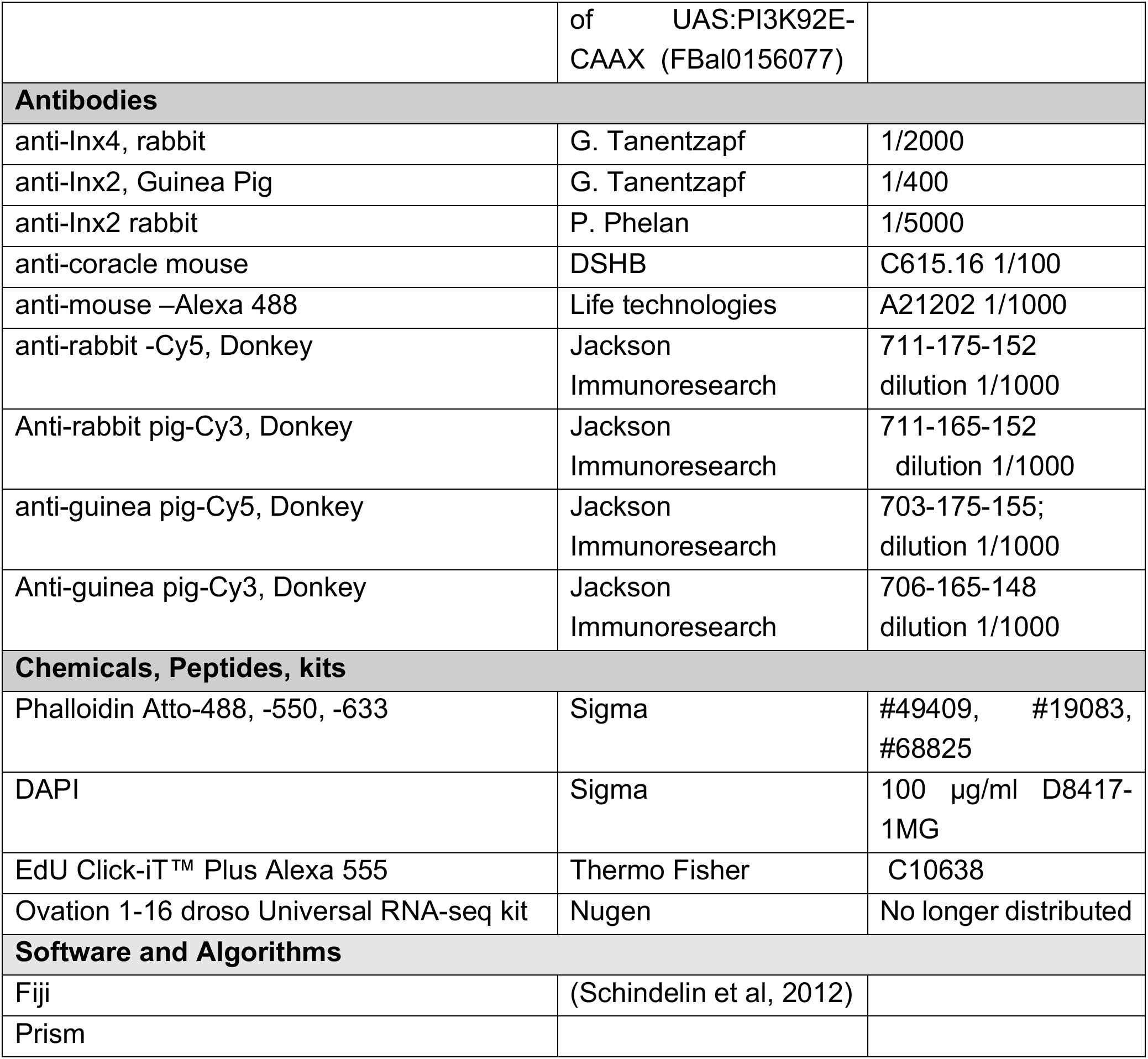
Resources and reagents.

**Table S3:**
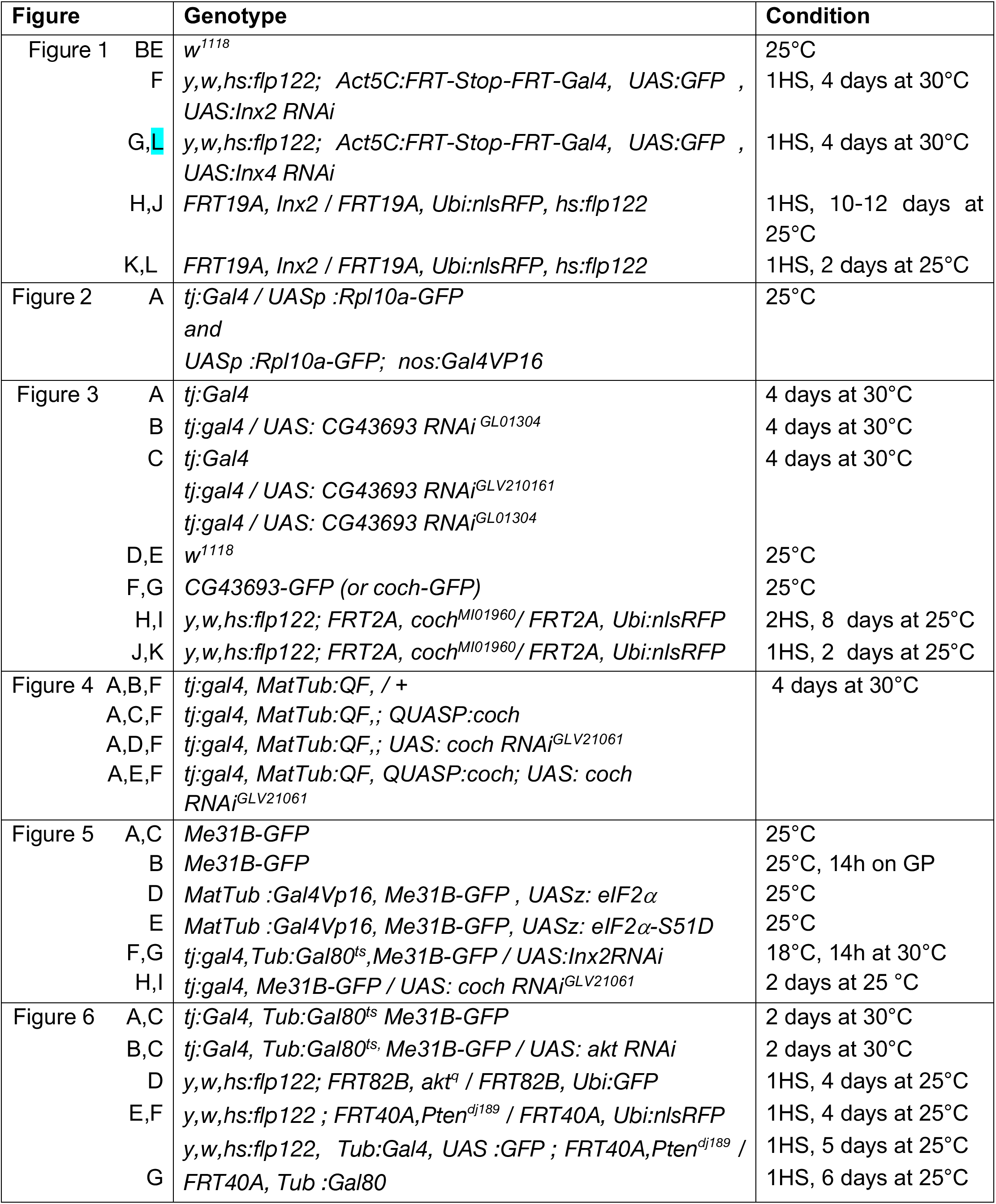
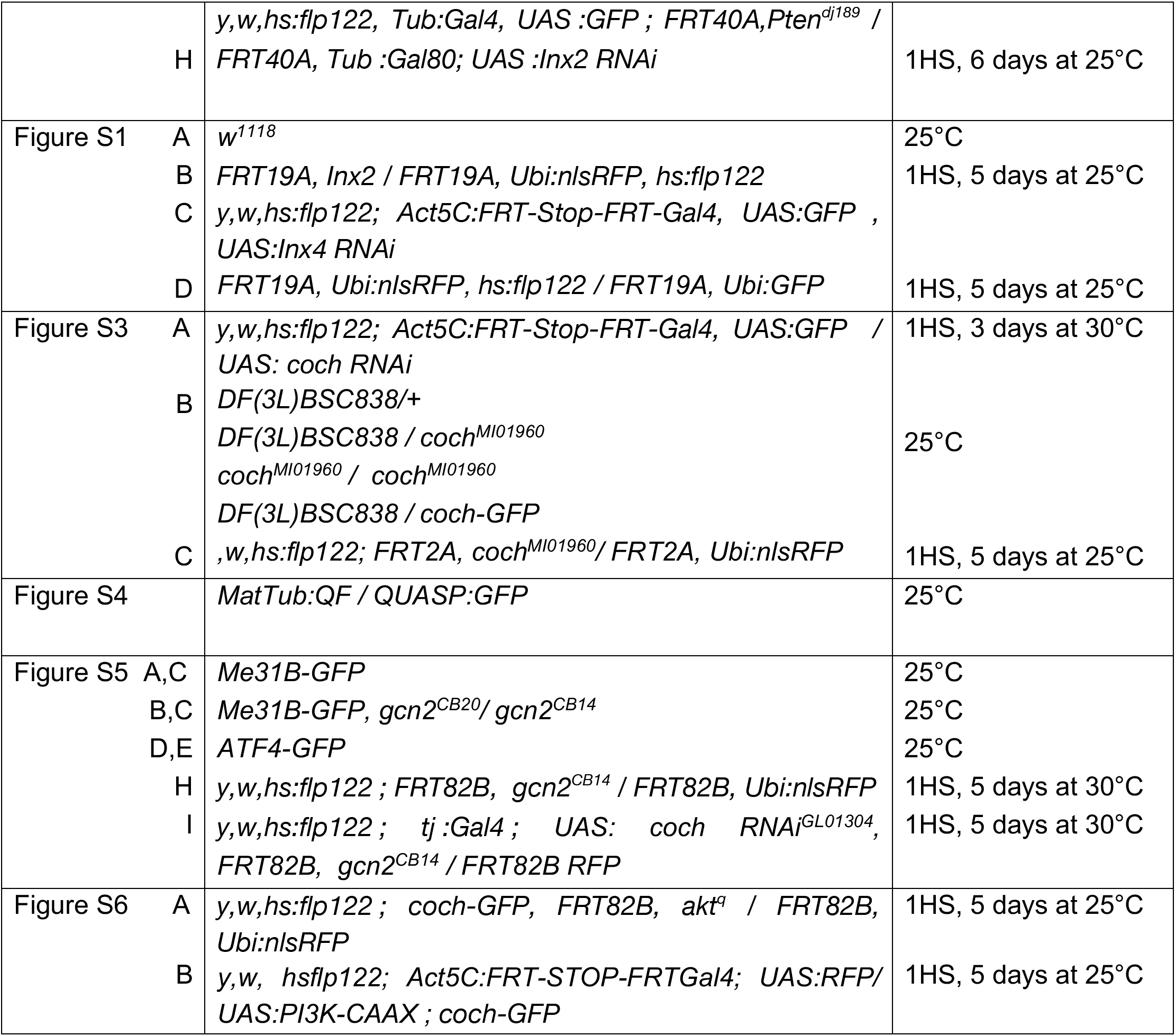
Genotypes and specific conditions. (h: hours; HS: heat-shock, GP : grape juice agar plate)

## Bibliography

1. Bauer R, Löer B, Ostrowski K, Martini J, Weimbs A, Lechner H, Hoch M. Intercellular communication: The drosophila innexin multiprotein family of gap junction proteins. Chemistry & Biology 2005, May;12(5).

2. Evans WH. Cell communication across gap junctions: A historical perspective and current developments. Biochem Soc Trans 2015, Jun;43(3).

3. Oshima A, Tani K, Fujiyoshi Y. Atomic structure of the innexin-6 gap junction channel determined by cryo-em. Nat Commun 2016;7.

4. Skerrett IM, Williams JB. A structural and functional comparison of gap junction channels composed of connexins and innexins. Dev Neurobiol 2017, May;77(5).

5. Elias LA, Wang DD, Kriegstein AR. Gap junction adhesion is necessary for radial migration in the neocortex. Nature 2007;448(7156).

6. Miao G, Godt D, Montell DJ. Integration of migratory cells into a new site in vivo requires channel-independent functions of innexins on microtubules. Dev Cell 2020;54(4).

7. Gittens JE, Kidder GM. Differential contributions of connexin37 and connexin43 to oogenesis revealed in chimeric reaggregated mouse ovaries. J Cell Sci 2005;118(Pt 21).

8. Simon AM, Goodenough DA, Li E, Paul DL. Female infertility in mice lacking connexin 37. Nature 1997;385(6616).

9. Colonna R, Mangia F. Mechanisms of amino acid uptake in cumulus-enclosed mouse oocytes. Biol Reprod 1983, May;28(4).

10. Haghighat N, LJ VW. Developmental change in follicular cell-enhanced amino acid uptake into mouse oocytes that depends on intact gap junctions and transport system gly. The Journal of Experimental Zoology 1990, Jan;253(1).

11. Eppig JJ, Pendola FL, Wigglesworth K, Pendola JK. Mouse oocytes regulate metabolic cooperativity between granulosa cells and oocytes: Amino acid transport. Biol Reprod 2005, Aug;73(2).

12. Sugiura K, Pendola FL, Eppig JJ. Oocyte control of metabolic cooperativity between oocytes and companion granulosa cells: Energy metabolism. Dev Biol 2005;279(1).

13. Doherty CA, Amargant F, Shvartsman SY, Duncan FE, Gavis ER. Bidirectional communication in oogenesis: A dynamic conversation in mice and drosophila. Trends Cell Biol 2022, Apr;32(4).

14. Church SH, Donoughe S, BAS DM, Extavour CG. Insect egg size and shape evolve with ecology but not developmental rate. Nature 2019, Jul;571(7763).

15. Petkova MD, Little SC, Liu F, Gregor T. Maternal origins of developmental reproducibility. Curr Biol 2014, Jun 2;24(11):1283–8.

16. Burn KM, Shimada Y, Ayers K, Vemuganti S, Lu F, Hudson AM, Cooley L. Somatic insulin signaling regulates a germline starvation response in drosophila egg chambers. Dev Biol 2015;398(2).

17. Doherty CA, Diegmiller R, Kapasiawala M, Gavis ER, Shvartsman SY. Coupled oscillators coordinate collective germline growth. Dev Cell 2021;56(6).

18. Vachias C, Fritsch C, Pouchin P, Bardot O, Mirouse V. Tight coordination of growth and differentiation between germline and soma provides robustness for drosophila egg development. Cell Rep 2014, Oct 23;9(2):531–41.

19. Spradling AC, Bate M, Arias AM. The development of drosophila melanogaster. The Development of Drosophila Melanogaster 1993;1:1–70.

20. Chen DY, Crest J, Streichan SJ, Bilder D. Extracellular matrix stiffness cues junctional remodeling for 3D tissue elongation. Nat Commun 2019;10(1).

21. Drummond-Barbosa D, Spradling AC. Stem cells and their progeny respond to nutritional changes during drosophila oogenesis. Dev Biol 2001, Mar 1;231(1):265–78.

22. LaFever L, Feoktistov A, Hsu HJ, Drummond-Barbosa D. Specific roles of target of rapamycin in the control of stem cells and their progeny in the drosophila ovary. Development 2010, Jul;137(13):2117–26.

23. Sun P, Quan Z, Zhang B, Wu T, Xi R. TSC1/2 tumour suppressor complex maintains drosophila germline stem cells by preventing differentiation. Development 2010, Aug 1;137(15):2461–9.

24. Maines JZ, Stevens LM, Tong X, Stein D. Drosophila dmyc is required for ovary cell growth and endoreplication. Development 2004, Feb;131(4):775–86.

25. Wang Y, Riechmann V. The role of the actomyosin cytoskeleton in coordination of tissue growth during drosophila oogenesis. Curr Biol 2007, Aug 7;17(15):1349–55.

26. Borreguero-Muñoz N, Fletcher GC, Aguilar-Aragon M, Elbediwy A, Vincent-Mistiaen ZI, Thompson BJ. The hippo pathway integrates pi3k-akt signals with mechanical and polarity cues to control tissue growth. PLoS Biol 2019, Oct;17(10):e3000509.

27. Tazuke SI, Schulz C, Gilboa L, Fogarty M, Mahowald AP, Guichet A, et al. A germline-specific gap junction protein required for survival of differentiating early germ cells. Development 2002, May;129(10).

28. Bohrmann J, Zimmermann J. Gap junctions in the ovary of drosophila melanogaster: Localization of innexins 1, 2, 3 and 4 and evidence for intercellular communication via innexin-2 containing channels. BMC Dev Biol 2008;8:111.

29. Sahu A, Karmakar S, Halder S, Ghosh G, Acharjee S, Dasgupta P, et al. Germline soma communication mediated by gap junction proteins regulates epithelial morphogenesis. PLoS Genet 2021;17(8).

30. Gilboa L, Forbes A, Tazuke SI, Fuller MT, Lehmann R. Germ line stem cell differentiation in drosophila requires gap junctions and proceeds via an intermediate state. Development 2003, Dec;130(26).

31. Mukai M, Kato H, Hira S, Nakamura K, Kita H, Kobayashi S. Innexin2 gap junctions in somatic support cells are required for cyst formation and for egg chamber formation in drosophila. Mech Dev 2011, Sep;128(7-10).

32. Pesch YY, Dang V, Fairchild MJ, Islam F, Camp D, Kaur P, et al. Gap junctions mediate discrete regulatory steps during fly spermatogenesis. PLoS Genet 2022;18(9).

33. Smendziuk CM, Messenberg A, Vogl AW, Tanentzapf G. Bi-directional gap junction-mediated soma-germline communication is essential for spermatogenesis. Development 2015;142(15).

34. Vachias C, Fritsch C, Pouchin P, Bardot O, Mirouse V. Tight coordination of growth and differentiation between germline and soma provides robustness for drosophila egg development. Cell Rep 2014, Oct 23;9(2):531–41.

35. Domanitskaya E, Anllo L, Schüpbach T. Phantom, a cytochrome P450 enzyme essential for ecdysone biosynthesis, plays a critical role in the control of border cell migration in drosophila. Dev Biol 2014;386(2).

36. Venken KJ, Schulze KL, Haelterman NA, Pan H, He Y, Evans-Holm M, et al. MiMIC: A highly versatile transposon insertion resource for engineering drosophila melanogaster genes. Nat Methods 2011, Sep;8(9):737–43.

37. Isasti-Sanchez J, Münz-Zeise F, Lancino M, Luschnig S. Transient opening of tricellular vertices controls paracellular transport through the follicle epithelium during drosophila oogenesis. Dev Cell 2021;56(8).

38. Potter CJ, Tasic B, Russler EV, Liang L, Luo L. The Q system: A repressible binary system for transgene expression, lineage tracing, and mosaic analysis. Cell 2010;141(3).

39. Rørth P. Gal4 in the drosophila female germline. Mech Dev 1998, Nov;78(1-2).

40. Shimada Y, Burn KM, Niwa R, Cooley L. Reversible response of protein localization and microtubule organization to nutrient stress during drosophila early oogenesis. Dev Biol 2011;355(2).

41. Riggs CL, Kedersha N, Ivanov P, Anderson P. Mammalian stress granules and P bodies at a glance. J Cell Sci 2020;133(16).

42. Cadena Sandoval M., Heberle AM, Rehbein U, Barile C, JM RP, Thedieck K. MTORC1 crosstalk with stress granules in aging and age-related diseases. Frontiers in Aging 2021;2.

43. Kosakamoto H, Okamoto N, Aikawa H, Sugiura Y, Suematsu M, Niwa R, et al. Sensing of the non-essential amino acid tyrosine governs the response to protein restriction in drosophila. Nat Metab 2022, Jul;4(7).

44. Srivastava A, Lu J, Gadalla DS, Hendrich O, Grönke S, Partridge L. The role of GCN2 kinase in mediating the effects of amino acids on longevity and feeding behaviour in drosophila. Frontiers in Aging 2022;3.

45. Kang MJ, Vasudevan D, Kang K, Kim K, Park JE, Zhang N, et al. 4E-BP is a target of the GCN2-ATF4 pathway during drosophila development and aging. J Cell Biol 2017;216(1).

46. Jewell JL, Russell RC, Guan KL. Amino acid signalling upstream of mtor. Nat Rev Mol Cell Biol 2013, Mar;14(3).

47. Carvalho-Santos Z, Cardoso-Figueiredo R, Elias AP, Tastekin I, Baltazar C, Ribeiro C. Cellular metabolic reprogramming controls sugar appetite in drosophila. Nat Metab 2020, Sep;2(9).

48. Su YQ, Sugiura K, Eppig JJ. Mouse oocyte control of granulosa cell development and function: Paracrine regulation of cumulus cell metabolism. Semin Reprod Med 2009, Jan;27(1):32–42.

49. Rodríguez-Nuevo A, Torres-Sanchez A, Duran JM, De Guirior C, Martínez-Zamora MA, Böke E. Oocytes maintain ros-free mitochondrial metabolism by suppressing complex I. Nature 2022, Jul;607(7920):756–61.

50. Cerqueira Campos F, Dennis C, Alégot H, Fritsch C, Isabella A, Pouchin P, et al. Oriented basement membrane fibrils provide a memory for f-actin planar polarization via the dystrophin-dystroglycan complex during tissue elongation. Development 2020;147(7).

51. Aigouy B, Umetsu D, Eaton S. Segmentation and quantitative analysis of epithelial tissues. Methods Mol Biol 2016;1478:227–39.

52. Aigouy B, Mirouse V. ScientiFig: A tool to build publication-ready scientific figures. Nat Methods 2013, Oct 30;10(11):1048.

